# Comparative analyses reveal rapid turnover and emergence of transitory 3D genome architectures in the fungal kingdom

**DOI:** 10.64898/2026.07.21.739809

**Authors:** Alice Laigle, Camille Cornet, Ashwini V. Mohan, Tobias Baril, Daniel Croll

**Affiliations:** Laboratory of Evolutionary Genetics, Institute of Biology, University of Neuchâtel, Neuchâtel, Switzerland; Biodiversity Genomics Laboratory, Institute of Biology, University of Neuchâtel, Neuchâtel, Switzerland

**Author notes:** Corresponding author:* Daniel Croll.

**Keywords:** chromatin folding, Fungi, Hi-C, comparative genomics, transposable elements, genomic interactions, biodiversity

## Abstract

The three-dimensional architecture of genomes plays major roles in biological processes such as gene expression and DNA replication. The architecture of genomes has evolved substantially with distinct 3D genome shapes being identified in different lineages. The factors driving the evolution of genome architectures have primarily been assessed in animals and plants, yet large parts of the tree of life remain poorly explored. Fungi offer excellent models to assess the evolution of 3D genome architecture in a phylogenetic context given rapid genome size changes and chromosomal sequence turnover. Here, we analyzed chromosome conformation data (Hi-C) of 55 fungal species with completely assembled genomes. We identified ten species with Rabl, one species with chromosome territories and ten with a novel, intermediate chromosomal architecture, where centromeres and telomeres are at opposites in the nucleus (Rabl-like) but with a distinct 3D organization. This “bean” shape likely evolved several times independently. The discovery of a genome with a chromosome territories conformation was unexpected, as this was thought to be associated with condensin II subunits in the animal kingdom. We investigated whether 3D conformations correlated with genome size and repeat content using phylogenetic independent contrasts, however we found no genomic feature to be significantly associated with changes in genome architecture. Overall, we report the first large-scale comparison of 3D genome architecture in the fungal kingdom and identify a novel “bean” configuration.

**Significance:** Three-dimensional genome architecture strongly influences gene regulation, yet little is known about 3D genome architecture in an organismal group that has adapted to nearly all ecosystems on our planet, Fungi. We reconstructed 3D genome architectures from 55 fungal species covering three different phyla and demonstrate that most species do not conform to the existing definitions of 3D architectures. We identified the first case of ‘Chromosome Territories’ in the Fungal Kingdom and a previously undescribed organization that we label "bean-shaped", and show that some fungal species do not conform to the canonical 3D-architecture categories of the animal and plant kingdoms. The diversity of genome architectures observed in the study could reflect the diverse gene regulatory mechanisms known from Fungi and marks the beginning of mapping out 3D genome organizations in this diverse clade. Further research in this area will uncover the diverse strategies employed by Fungi in light of their rapid adaptation.

## Introduction

The three-dimensional (3D) genome architecture plays a crucial role in facilitating physical interactions that regulate gene expression during key biological processes (Kumar *et al.,* 2021; Tavallaee and Orouji, 2025). These processes encompass development and cell differentiation (Dixon *et al.,* 2015; Camerino *et al.,* 2024), adaptations to biotic and abiotic stress (Sun *et al.,* 2020; Huang *et al.,* 2023; Marie *et al.,* 2024), and speciation (Yamasaki *et al.,* 2025). The 3D architecture of genomes facilitates physical interactions between *cis*-regulatory elements and their target genes (Tavallaee and Orouji, 2025), as well as promoter-to-promoter and enhancer-to-promoter interactions (Hsieh *et al.,* 2020; Papantonis and Oudelaar, 2025). Disruption of this 3D genome architecture can result in dysfunctional gene expression, which has been linked to various diseases, including cancer (Li *et al.,* 2018; Tettey *et al.,* 2023). Yamasaki *et al.* (2025) showed that 3D architecture facilitates the establishment of barriers to gene flow and therefore could have an impact on speciation. Yet, our understanding of 3D genome architecture and its consequences remains limited to narrow phylogenetic groups and is not yet broadly generalisable due to insufficient sampling across the tree of life and ongoing methodological developments. An expanded perspective on 3D genome architectures across the tree of life will provide insights into evolutionary processes leading to innovations in chromosomal organization.

During interphase, the 3D genome architectures that have been identified fall in one of the two primary shape categories depending on the species: Rabl and chromosome territories (Hoencamp *et al.,* 2021). In the Rabl organization (also called type I), telomeres and centromeres are polarized inside the nucleus (Rabl, 1885; Hoencamp *et al.,* 2021). The Rabl organization has been subdivided into telomere clustering, centromere-telomere axis, or centromere clustering (Hoencamp *et al.,* 2021), which will all three be referred to as Rabl-like. In the chromosome territories organization (also called type II), chromosomes occupy spatially defined locations in the nucleus, with no opposite positioning of telomeres and centromeres (Boveri, 1888; Cremer and Cremer, 2001). Generally, vertebrates exhibit chromosome territories (Cremer and Cremer, 2001; Lieberman-Aiden *et al.,* 2009; Li *et al.,* 2022), while species lacking condensin II subunits, such as the fruit fly *Drosophila melanogaster* and the moss animal *Cristatella mucedo* (Hoencamp *et al.,* 2021), present type I organizations. During interphase, plants mostly exhibit type I organization, as observed in bread wheat and the ground peanut (Concia *et al.,* 2020; Hoencamp *et al.,* 2021). Most studies on 3D genome architectures to date have focused on animals and plants, whilst 3D genome architectures in other eukaryotes, including Fungi, remain poorly understood.

3D genome architectures investigated in Fungi to date follow Rabl (type I) organizations including model yeasts and rusts from the genus *Puccinia* (Tanizawa *et al.,* 2010; Belton *et al.,* 2015; Xia *et al.,* 2022). The arbuscular mycorrhizal symbiont *Rhizophagus irregularis* presents a possible exception with an intermediate organization between Rabl and chromosome territories (Yildirir *et al.,* 2022). Fungi with a centromere clustering organization include *Neurospora crassa* and *Epichloë festucae*, (Galazka *et al.,* 2016; Winter *et al.,* 2018). In the human pathogen *Candida albicans*, Burrack *et al.* (2016) demonstrated that if primary centromeres cluster together, neo-centromeres (*i.e.* new centromeres formed in a region without centromeric sequences) cluster as well and are functional. Some plant pathogens share an organization where core chromosomes display centromere clustering, while accessory chromosomes (*i.e.* chromosomes not shared by all members of the same species) are aggregated together (Wang *et al.,* 2024; Glavincheska and Lorrain, 2025). Given limited macroevolutionary perspectives on the evolution of 3D genome architectures, comparative analyses of chromosomal conformation are needed to identify the diversity, prevalence, and functional impacts of genome configurations across major fungal lineages. High-throughput sequencing coupled with chromosome conformation capture (Hi-C; Lieberman-Aiden *et al.,* 2009) provides an efficient approach to systematically capture interactions along the genome and to reconstruct chromosomal conformations for a population of cells.

In this study, we investigated publicly available Hi-C datasets generated by the Darwin Tree of Life project (DToL; The Darwin Tree of Life Project Consortium, 2022) using a standardized pipeline to investigate diversity in 3D genome architectures in a phylogenetic context. This dataset encompasses a total of 54 species spanning three different phyla for which data from a Hi-C protocol was available. In addition, we re-analyzed an already published dataset of *R. irregularis* as we only had one species available in the Mucoromycota phylum (Yildirir *et al.,* 2022). By characterizing 3D genome architecture among species, we assess the diversity of genomic architectures and investigate whether genomic features such as repeat content could account for changes in 3D structures.

## Results

### Diversity of 3D genome architectures

To explore 3D genome architecture diversity in the fungal kingdom, we collected publicly available Hi-C data with their corresponding chromosome-level assembled genomes. The DToL project offered the most homogenous dataset, characterized by consistent protocols for genome assemblies and Hi-C data generation (The Darwin Tree of Life Project Consortium, 2022; Supplemental Table 1). We omitted datasets showing less than one million valid pairs after duplicates removal (see Methods; Supplemental Table 2). We retained a total of 54 species from three phyla: 45 Basidiomycota, 13 Ascomycota and a single Mucoromycota species (Supplemental Table 1). Considerable lifestyle diversity is present among the examined species, ranging from pathogenic rusts to symbiotic organisms such as mycorrhizal fungi and lichens, as well as a marine yeast and common saprophytic fungi. The sampling scope enabled us to assess genome architecture considering both genetic and ecological factors. Additionally, we incorporated the *R. irregularis* DAOM-197198 strain (Yildirir *et al.,* 2022) for comparison with *Entomortierella parvispora*, the only Mucoromycota species currently represented in the DToL project (Supplemental Table 1). We constructed a supermatrix phylogenetic tree based on BUSCO genes from all the species in the dataset (see Methods). In addition, we included three outgroup species (*Conidiobolus obscurus*, *Coemansia mojavensis*, *Linderina pennispora*) from the Zoopagomycota phylum. The maximum likelihood tree is robust, with only three nodes showing bootstrap scores below 100 (*i.e.*, 84-96; Figure 1A).

**Figure 1.**
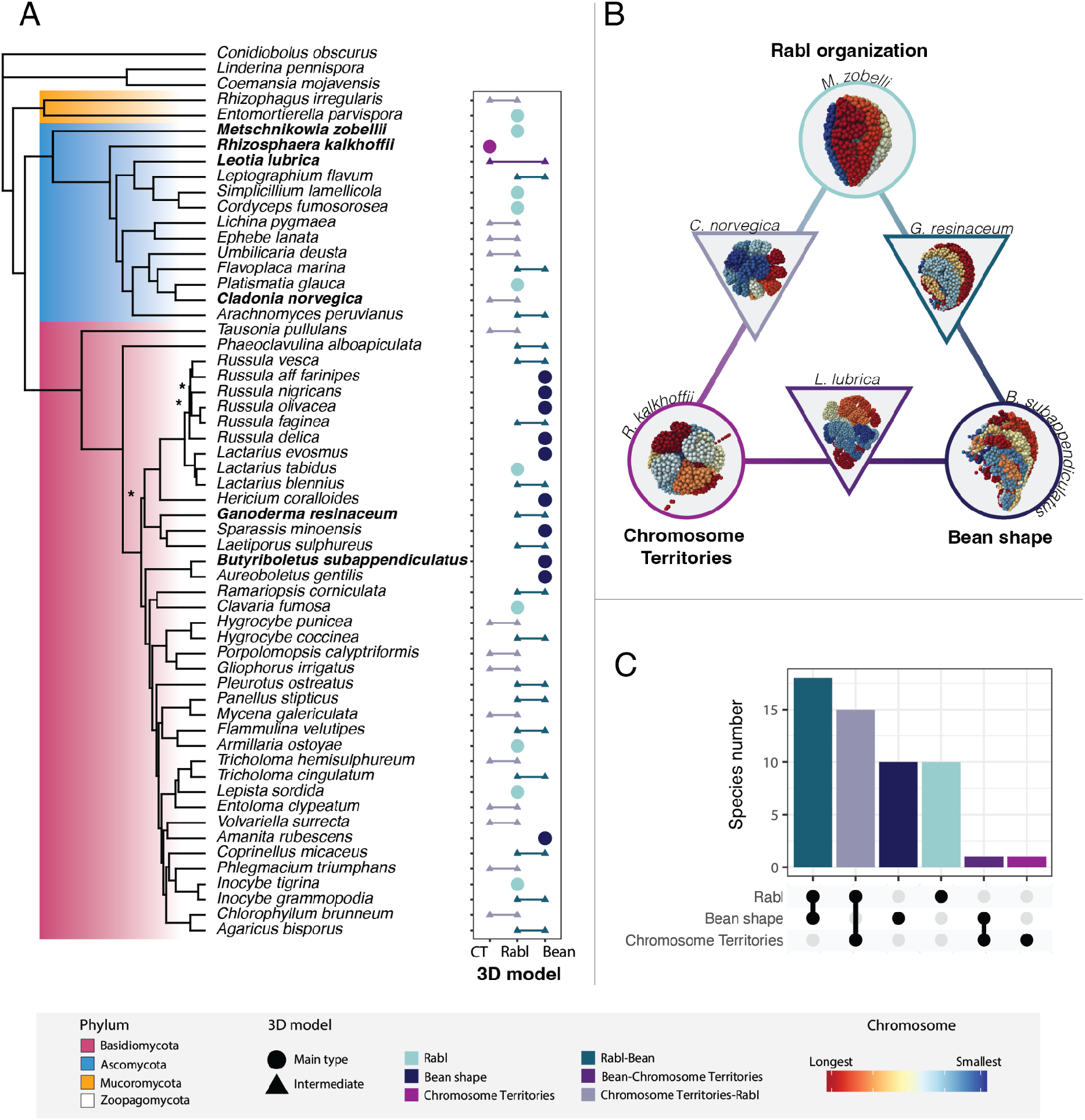
Diversity in 3D genome architectures among Fungi. **A.** Maximum likelihood phylogenetic tree of the 55 analyzed species and three outgroups (*Conidiobolus obscurus*, *Coemansia mojavensis*, *Linderina pennispora*) with a dot plot representing the 3D genome architectures. Colored backgrounds of the tree highlight the phyla: Zoopagomycota (white: outgroup), Mucrocomycota (orange), Ascomycota (blue) and Basidiomycota (pink). Bold species names refer to the examples given in (B). Asterisks in the tree represent bootstrap support values of 84-86. The customized dot plot shows the 3D models of each species. Circles are for the main 3D genome architectures: Rabl organization, chromosome territories (CT) and Bean shape. Linked triangles represent intermediate 3D genome architectures found in the dataset: Rabl-Bean, Bean-chromosome territories and chromosome territories-Rabl. Colors of circles and triangles represent the 3D models: Rabl (light green), bean (dark blue), chromosome territories (fuschia), Rabl-bean (duck green), Bean-chromosome territories (purple) and chromosome territories-Rabl (clear purple). **B.** Representative 3D genome architectures discovered among the analyzed species. The triangular gradient represents discoveries of intermediate forms of 3D genome architecture between primary shapes. Shapes and colors follow the color code as in **A**. Each bead represents 10 kb of a chromosome and is colored as a gradient to represent the chromosome it belongs to, from red for the longest chromosome to dark blue for the smallest one. The complete set of 3D models and related contact maps are shown in Supplemental Figure 1 and rotating 3D models are available on Zenodo (10.5281/zenodo.17672270). **C.** Upset plot summarizing the number of species displaying the different 3D genome architectures and their intermediates.

We characterized the 3D genome architecture of each species with high-quality Hi-C datasets by categorizing them according to previously identified genome architectures (*i.e.,* Rabl and chromosome territories). Upon visual inspection, we discovered that several genomes displayed an intermediate form of organization, which we describe as bean-shaped. In the bean organization, most centromeres cluster together and chromosomal arms stretch in a curved way (Figure 1B; Supplemental Figure 1). Bean shapes were only found in the Basidiomycota phylum (Figure 1A; Supplemental Figure 1). Beyond the bean-like shape, we found that most species fall into a gradient of intermediate forms between the extremes of Rabl and chromosome territories. To further investigate this, we defined a gradient of organizational types with bean shape as one of the main categories, alongside Rabl and chromosome territories (Figure 1B). Among the main organizational types, ten species displayed a Rabl organization, and another ten species exhibited a bean shape. Notably, only *Rhizosphaera kalkhoffii* (Ascomycota) demonstrated clearly defined chromosome territories (Figure 1; Supplemental Figure 1). Among the intermediates, fifteen species were categorized as chromosome territories-Rabl intermediate, thirteen species as Rabl-bean intermediate, and only *Leotia lubrica* (Ascomycota) as bean-chromosome territories intermediate (Figure 1; Supplemental Figure 1). Among the species exhibiting Rabl-bean intermediates, several displayed a distinct pattern in the Hi-C contact maps, which we characterized as a compact bean shape (*e.g., Ganoderma resinaceum*; Supplemental Figure 1). In these maps, the diagonal contacts converge at specific chromosomal locations, such as in *G. resinaceum*, *Phaeoclavulina alboapiculata*, and *Pleurotus ostreatus* (Figure 1B; Supplemental Figure 1). Overall, we found more 3D genome architecture types in Ascomycota compared to Basidiomycota. Interestingly, within Basidiomycota, the bean organization appears to have multiple independent origins (Figure 1), however this needs to be re-evaluated with a wider phylogenetic sampling and the inference of ancestral genome architecture states.

### 3D architectures and genome characteristics

We investigated genomic characteristics including genome size, gene and repeat content to assess whether these were correlated with 3D genomic architectures. Genome sizes ranged from 13.6 to 165.0 Mbp, with the marine yeast *Metschnikowia zobellii* (Ascomycota) and the mycorrhizal symbiont *Phlegmacium triumphans* (Basidiomycota) representing the smallest and largest genomes, respectively (Figure 2; Supplemental Table 3). Genome size was found to be significantly greater among the included Basidiomycota species compared to the represented Ascomycota species (Welch two-sample *t*-test: *t_33.7_* = −2.6, *p*-value = 0.013; Supplemental Figure 2). The GC content varied from 27.84 % in the arbuscular mycorrhizal symbiont *R. irregularis* (Mucoromycota) to 59.92 % in the phytopathogen *Leptographium flavum* (Ascomycota), with a median GC content of 49.43 % (Figure 2; Supplemental Table 3). Additionally, the number of nuclear chromosomes ranged from 5 to 32, corresponding to *M. zobellii* and *R. irregularis*, respectively, with a median of 13 chromosomes (Figure 2; Supplemental Table 3). Following curation and annotation of transposable elements (TEs) (Supplemental Table 4), we examined the distribution of gene and TE content across genomes in 10 kb bins using a subset of species with high-resolution Hi-C data (Supplemental Figure 3; Supplemental Table 3). Gene content varied significantly, ranging from 7.9% in *Phlegmacium triumphans* to 56.1% in the yeast *Tausonia pullulans* (Basidiomycota), with a median of 33.0%. Among the analyzed species, Ascomycota generally had a higher gene content than Basidiomycota species (Welch two-sample *t*-test: *t_19.9_* = 3.5, *p*-value = 2.1e^-3^; Supplemental Figure 2). Notably, the six species belonging to the Russulaceae family (genera *Lactarius* and *Russula*) exhibit high gene content with the maximum being 23.7 % in *Russula olivacea*. Among these, more than half of the Russulaceae species (5 out of 9) display bean shape organization, which also accounts for half of all species exhibiting this configuration among the analyzed species (5 out of 10; Supplemental Tables 3 and 5). TE content exhibited even greater variability than gene content, ranging from 1.4 % in the endo-parasitic fungus *Simplicillium lamellicola* to 78.8 % in the slimy waxcap *Gliophorus irrigatus*, but with a similar median of 31.1 % (Figure 2; Supplemental Figure 3; Supplemental Table 3). Among the included species, Basidiomycota possessed significantly higher TE content compared to Ascomycota (Welch two-sample *t*-test: *t_19.9_* = −3.1, *p*-value = 0.005; Supplemental Figure 2). We further investigated TEs classes subdivided into Class I retrotransposons, comprising Long Terminal Repeat (LTR) elements, Long Interspersed Nuclear Elements (LINE), and *Penelope*-Like Elements (PLE), and Class II elements, including DNA transposons and Rolling-Circle (RC) elements (also termed *Helitrons*) (Supplemental Table 6). Notably, the Short Interspersed Nuclear Elements (SINE, Class I) and those annotated broadly as retrotransposons were present at low percentages across species (Figure 2; Supplemental Table 6). We categorized the few retroposons under LTRs. Despite higher TE content in Basidiomycota, we did not observe any significant differences between the Ascomycota and Basidiomycota phyla in any of the TE categories mentioned above. However, it is noteworthy that *Ramariopsis corniculate* (Basidiomycota) displayed a high proportion of RC elements, accounting for 22.6 % of its genome (Figure 2; Supplemental Table 6).

**Figure 2.**
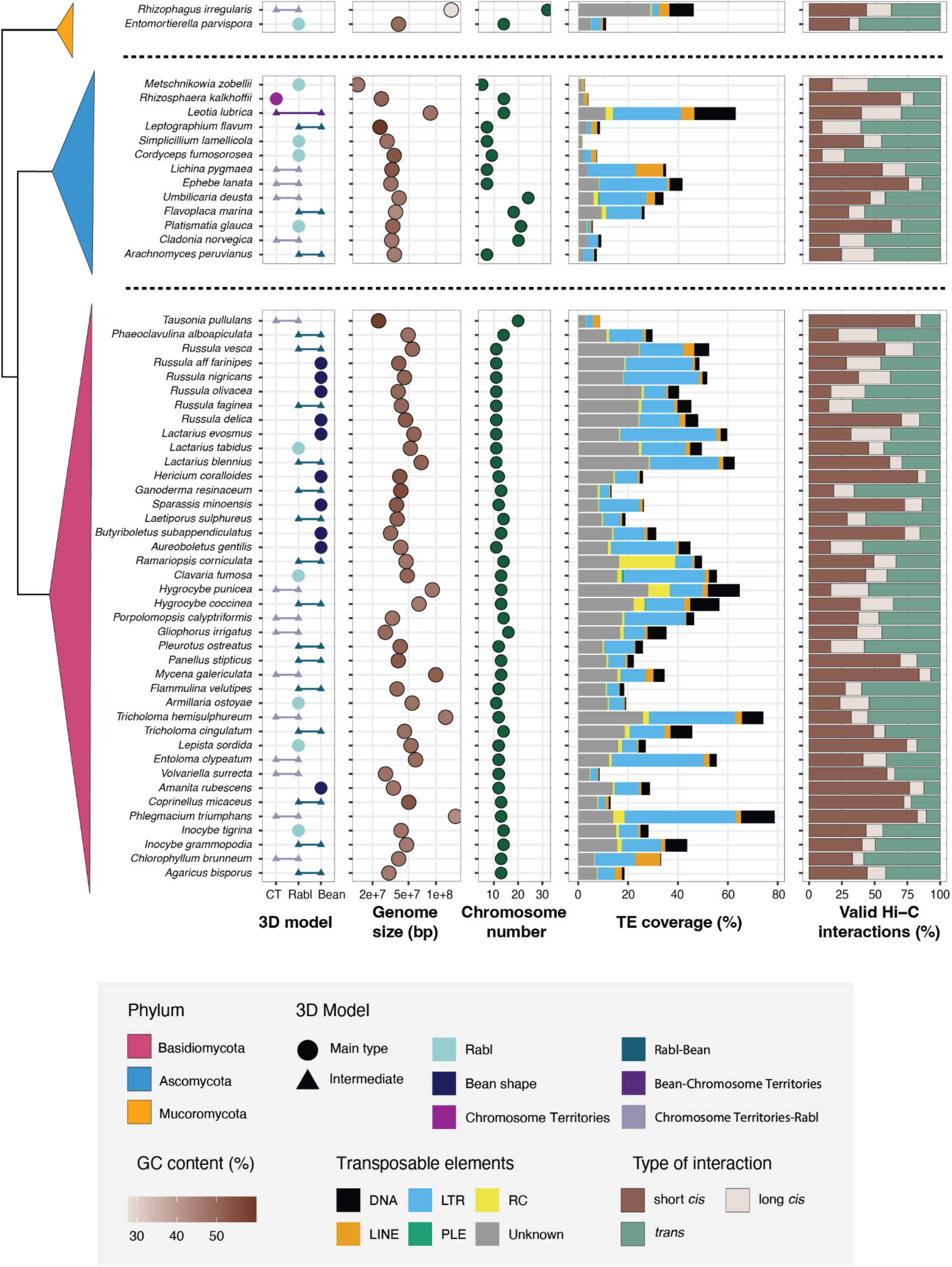
Associations between genomic features and 3D genome architectures. From left to right: **1)** schematic representation of the phylogenetic tree for the 55 species to distinguish their respective phylum. **2)** Dot plot representing the 3D genome architectures as presented in Figure 1. **3)** Dot plot showing the genome size (min: 13.60 Mb, max: 164.98 Mb, median: 39.94 Mb) colored by their GC content (min: 27.84, max: 59.92, median: 49.43). **4)** Dot plot of the number of chromosomes for each species (min: 5, max: 32, median: 13). **5)** Stack bar plot of the transposable elements (min: 1.44 %, max: 78.76 %, median: 31.11 %). SINE elements occurred at less than 0.2 % percent and are not shown. **5)** 100 % stack bar plot of the valid Hi-C interactions from corrected iced matrices at 50 kb resolution. Short *cis* interactions (brown) represent all bins interacting with themselves or the bin next to it, long *cis* interactions (beige) being all the other interactions on the same chromosome. *Trans* interactions (green) are the interactions between bins on different chromosomes. Basic stats of *cis* interactions: min: 26.9 %, max: 92.7 %, median: 57.9 %; of *trans* interactions: min: 7.3 %, max: 73.1 %, median: 42.1 %.

### 3D genome architectures are not associated with genomic features

To explore potential associations between genomic features and 3D genome architectures, we employed phylogenetic regressions and phylogeny-informed ANOVAs (Figure 3; Supplemental Figure 4; Supplemental Table 7). We first assessed whether the number of interactions reported by the Hi-C analyses was correlated with genome size and found no significant relationship (PGLS: *t_55_* = 0.91; *p*-value = 0.365; Supplemental Table 7). Hence, we expect that differences in genome size would not bias contact frequency landscapes across the studied species (Supplemental Figure 4A; Supplemental Table 7). We then investigated whether various genomic features were associated with each other. As expected, genome size showed a strong positive association with TE content (PGLS: *t_55_* = 7.79; *p*-value = 2.5e^-10^; Figure 3A; Supplemental Table 7). Additionally, genome size was positively associated with chromosome number (PGLS: *t_55_* = 2.23; *p*-value = 0.030) but negatively associated with gene content (PGLS: *t_55_* = −9.76; *p*-value = 1.9e^-13^) and GC content (PGLS: *t_55_* = 5.72; *p*-value = 3.43e^-05^; Supplemental Figure 4B-D; Supplemental Table 7). Furthermore, GC content was also negatively associated with TE content (PGLS: *t_55_* = −3.31; *p*-value = 1.7e^-3^; Supplemental Figure 4E). We then examined whether specific TE types were associated with genome size. We discovered that both Class I LTR and Class II DNA elements exhibited strong positive associations with genome size (PGLS: *t_55_* = 4.54; *p*-value = 3.23e^-05^ and *t_55_* = 6.83; *p*-value = 8.45e^-09^, respectively; Supplemental Figure 4F-G; Supplemental Table 7), but neither Class I LINE, Class I PLE nor Class II RC elements were associated with genome size (Supplemental Figure 4H-J; Supplemental Table 7). Interestingly, Class I SINE elements, ranging from 0 to 0.133% of the genome content, were also positively associated with genome size (PGLS: *t_55_* = 5.02; *p*-value = 6.25e^-06^; Supplemental Figure 4K; Supplemental Table 7).

**Figure 3.**
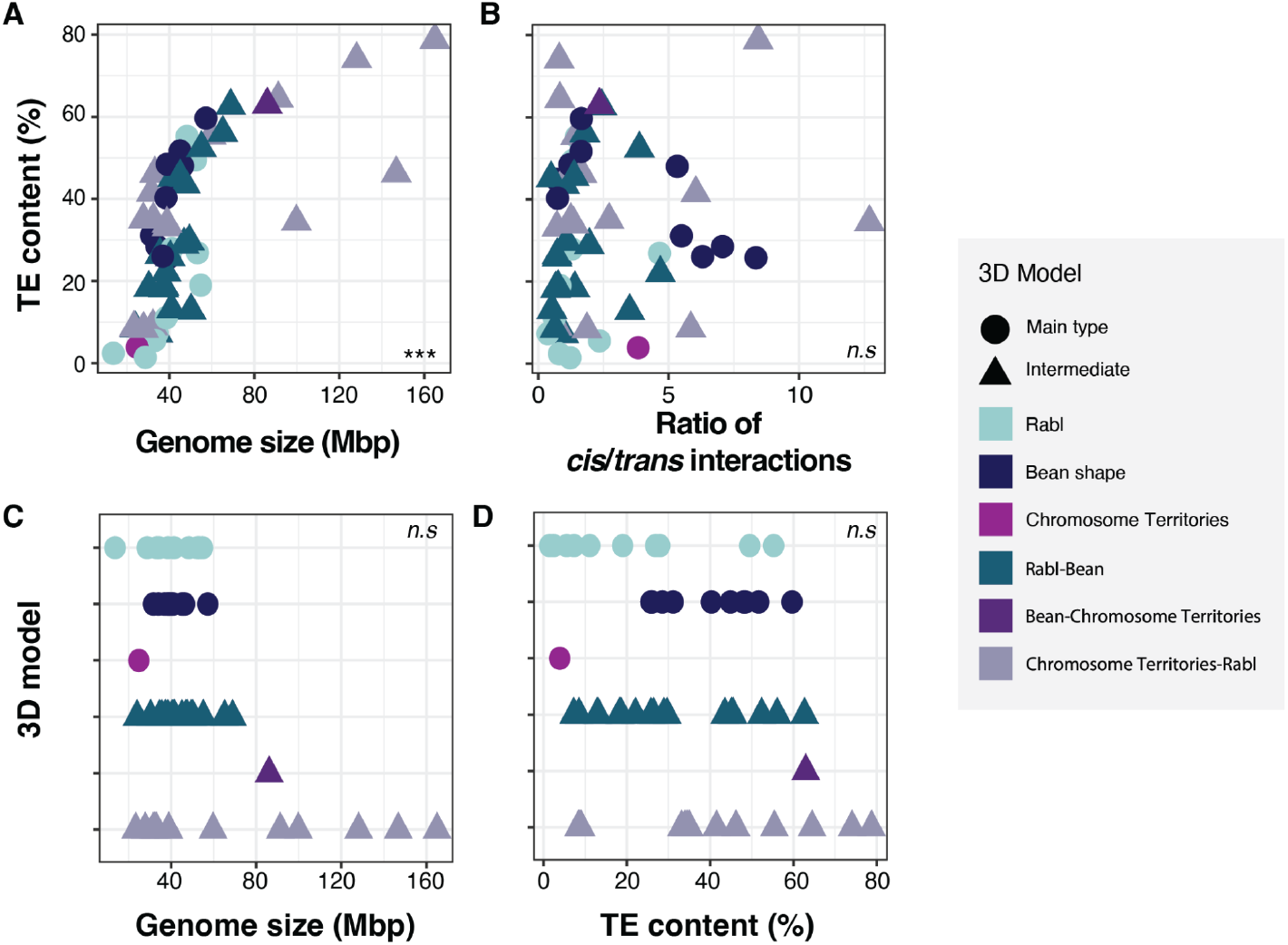
Associations between genetic features and 3D genome architecture. **A.** Dot plot illustrating the phylogenetic regression between genome size (Mbp) and TE content (%; *p*-value = 2.5e^-10^, AIC = 1953.41, BIC = 1962.29). **B.** Dot plot depicting the phylogenetic regression between the ratio of *cis*/*trans* interactions and TE content (%; *p*-value = 0.401, AIC = 271.18, BIC = 279.06). The ratio has been calculated by dividing the percentage of *cis* interactions by the percentage of *trans* interactions. **C.** Dot plot representing the phylogenetic ANOVA comparing the genome size (Mbp) and the 3D model (*p*-value = 0.336). **D.** Dot plot representing the phylogenetic ANOVA comparing TE content (%) and the 3D model (*p*-value = 0.166). Dot shapes and colors are as described in Figure 1.

Given that the genome structure affects *trans* interactions (defined here as interchromosomal interactions, in comparison to *cis* interactions which are intrachromosomal interactions; Duan *et al.,* 2010), we tested the association of the ratio *cis*/*trans* with TE content and genome size. We found that genome size and the *cis*/*trans* ratio were positively associated (PGLS: *t_55_* = 2.13; *p*-value = 0.038; Supplemental Figure 4L; Supplemental Table 7), but TE content and the *cis*/*trans* ratio were not associated (PGLS: *t_55_* = 0.85; *p*-value = 0.401; Figure 3B). Next, we examined associations between genome size and 3D genome architecture but found no significant correlation (phylogeny-informed ANOVA: *p*-value = 0.366; Figure 3C). Similarly, TE content and 3D genome architecture were not associated (phylogeny-informed ANOVA: *p*-value = 0.166; Figure 3D). Among all the other analyzed features, only Class II DNA elements were correlated with the 3D genome architecture (phylogeny-informed ANOVA: *p*-value = 0.025; Supplemental Figure 5F).

## Discussion

Here, we report a comparative investigation of 3D genome architectures in the fungal kingdom and identify a novel type of genome architecture called the bean organization. Genome architectures across the tree of life typically fall into two primary categories: Type I, which includes three Rabl-like organizations, and Type II, characterized by chromosome territories (Hoencamp *et al.,* 2021). We report a distinct organization that does not conform well to any established classification. The novel organization was found in Basidiomycota species and primarily in the better sampled Russulaceae family, specifically in the genera *Russula* and *Lactarius*. We further identified that this bean shape organization may have independently emerged multiple times within the Basidiomycota lineage. The novel organization may be associated with the plasticity of fungal nuclei. Characterisation of genome architectures in Fungi beyond those inferred using Hi-C approaches are rare, except for evidence for nuclear shapes in some Basidiomycota defined using transmission electron microscopy (Gao *et al.,*2019).

Our study also identified a species with chromosome territories organization, which is unexpected due to the loss of Condensin II in Fungi (Hoencamp *et al.* 2021). Hoencamp *et al.* (2021) proposed that Condensin II subunits are necessary in establishing chromosome territories. However, the observation of a genome architecture similar to those in mammals in another fungus, *R. kalkhoffii*, raises the question of how this organization arose despite the lack of Condensin II. A wider and denser sampling of fungal 3D genome architectures will be required to understand evolutionary transitions resulting in chromosome territories organization in Fungi. Beyond the discovery of novel and unexpected 3D architectures, we found that 3D genome architectures in Fungi are spread across gradients between the well-known Rabl and chromosome territories shapes, as well as the bean shape described here. This raises the question whether shape categories are truly discrete units. Increasing the phylogenetic breadth of species assessed for 3D architectures across the tree of life may well lead to further discoveries of genome architectures beyond the well-established categories. However, ascertaining new categories and intermediate shapes will require standardized Hi-C data and analyses. For instance, microscopy data alone would only reveal a spherical nucleus, obscuring the potential convergence of chromosomes where telomeres and centromeres may connect. Despite the considerable number of Hi-C datasets provided by the DToL, species representation across the fungal tree of life remains highly skewed and does not represent the phylogenetic breadth and species richness among major clades. For instance, our dataset is overrepresenting Basidiomycota compared to Ascomycota and lacks representation of pathogenic, early-diverging and unicellular fungal species.

Genome characteristics likely influence 3D architectures. We found a positive correlation between genome size and TE content as expected (Eliott and Gregory, 2015; Muszewska *et al.,* 2017; Oggenfuss *et al.,* 2021). However, we found no correlation between genome size and 3D genome architecture. This lack of correlation might be attributed to the uneven representation of the different architectural organizations, lack in phylogenetic breadth, and/or other factors governing genome architecture, for example epigenetic modifications. Interestingly, larger genomes were organized as intermediate shapes between chromosome territories and Rabl organizations indicating that genome size may constrain 3D genome architecture. We found that structures related to the bean shape and the intermediate between chromosome territories and Rabl organizations had a tendency towards higher ratio of *cis* to *trans* interactions. Conversely, Rabl organizations and their intermediates with bean shape showed an inverted interaction ratio.

This study demonstrates that 3D genome architecture is diverse across major lineages of the fungal kingdom, with the discovery of the bean organization. Intriguingly, many species display intermediate 3D organizations not fitting into the categories identified previously. Combined Hi-C generation and microscopy studies of nuclear organization across the fungal kingdom are necessary to unravel the evolutionary origins of 3D genome architectures such as novel shapes and convergent evolution of chromosome territories. Cataloguing the diversity of 3D genome architectures at scale across Fungi will provide an unprecedented opportunity to identify genomic drivers of chromosomal conformation.

## Materials and Methods

### Access of genome sequences and Hi-C datasets

All 54 fungal Hi-C data and assembled genomes used in this article are from the Darwin Tree of Life Project (October 2024-April 2025, https://portal.darwintreeoflife.org, The Darwin Tree of Life Project Consortium, 2022), except for *R. irregularis* strain DAOM-197198 (Yildirir *et al.,* 2022; Supplemental Table 1). Genomes were downloaded using the NCBI *datasets* method (*v16.20.1*; O’Leary *et al.,* 2024) and genome quality was inspected using QUAST (*v5.2.0*; Gurevich *et al.,* 2013; Supplemental Table 8). Chromosomes, contigs and scaffolds below 500 kb were removed using SAMtools *faidx* (*v1.21*; Danecek *et al.,* 2011) and seqtk *seq* methods (*v1.4*; Li, 2012). Hi-C data were downloaded from NCBI using *prefetch* and *fasterq-dump* functions from SRA-Tools (*v3.1.1*; https://github.com/ncbi/sra-tools; Sherry *et al.,* 2008). Three additional genomes from the Zoopagomycota phylum were downloaded and quality controlled as above to be used as outgroup species (*Coemansia mojavensis*, *Conidiobolus obscurus*, *Linderina pennispora*) for the phylogenetic tree construction (Supplemental Tables 1 and 7).

### Generation of the phylogenetic tree

To generate a rooted tree, three Zoopagomycota species were used as an outgroup (*Coemansia mojavensis*, *Conidiobolus obscurus* and *Linderina pennispora*). Universal single-copy genes were assessed for the 58 species using BUSCO with the *-l fungal_odb10* option (*v5.8.0*; Manni *et al.,* 2021) and only complete BUSCO genes were extracted. Among the complete BUSCO genes, only those found in at least three species were kept. Their amino acid sequences were then aligned using MAFFT (*v7.626*; Katoh and Standley, 2013). A supermatrix was produced with the *create_concat* of PhyKIT (*v2.0.1*; Steenwyk *et al.,* 2021) and ClipKIT (Steenwyk *et al.,* 2020). The tree was generated using IQ-TREE2 (*v2.3.6*; Minh *et al.,* 2020) with the ultrafast bootstrap option set to 1000 replicates (Hoang *et al.,* 2018), which invoked ModelFinder (Kalyaanamoorthy *et al.,* 2017). The best-fit model, according to BIC, was Q.insect+F+I+R10. The phylogenetic tree was forced ultrametric using the function *chronos()* of the package *ape* (*v5.8.1*) in R (Paradis *et al.,* 2004), and only three bootstraps were not at 100, ranging 84-86.

### Annotation of genomes

Gene annotations were made using the pipeline C of BRAKER2 (Hoff *et al.,* 2016; Bruna *et al.,* 2021; Hoff *et al.,* 2019; Bruna *et al.,* 2020; Lomsadze *et al.,* 2005; Buchfink *et al.,* 2015; Gotoh, 2008; Iwata and Gotoh, 2012; Ter-Hovhannisyan *et al.,* 2008; Stanke *et al.,* 2008; Stanke *et al.,* 2006), with the fungal OrthoDB 11 database (Kuznetsov *et al.,* 2022) and the *-- fungus* option. To annotate TEs, we processed as follows: constructed individual libraries using Earl Grey (Baril *et al.,* 2024) and created a single library for all studied species using CD-HIT (v4.8.1; Li and Godzik, 2006), curated it (Supplemental Table 4) and using SeqKit2 (v2.9.0; Shen *et al.,* 2024) and seqtk (v1.4-r132; https://github.com/lh3/seqtk; Li, 2012). We finally annotated genomes with this single curated library using *earlGreyAnnotationOnly* command from Earl Grey (Baril *et al.,* 2024).

### Treatment of Hi-C data, normalization and conversions

All Hi-C data were processed with the 3d-genome builder (3DGB) snakemake pipeline (*v1.1*, Poinsignon *et al.,* 2023) at 10, 20, 30, 50 and 100 kb resolutions. The pipeline integrates Bowtie2 (*v2.5.1*; Langmead and Salzberg, 2012), the HiC-Pro pipeline (*v3.1.0*; Servant *et al.,* 2015), the *iced* python package (Servant *et al.,* 2015), the *pastis* python package (Varoquaux *et al.,* 2014) and the *ggplot2* package (Wickham, 2016). Species having less than one million valid pairs after removing duplicates were discarded in the analyses (*Agrocybe praecox*, *Buglossoporus quercinus*, *Hericium erinaceus*, *Inocybe obscuroides*, *Ripartites tricholoma*; “valid_interaction_rmdup” in Supplemental Table 2). Raw matrices were converted to H5 file format at 50 kb resolution using *hicConvertFormat* from HiCExplorer (Ramírez *et al.,* 2018; Wolff *et al.,* 2020). *hicCorrectMatrix* with the *diagnostic_plot* option from HiCExplorer was used to define thresholds for matrix corrections before applying the thresholds with *hicCorrectMatrix* (Supplemental Table 9; Supplemental Figure 1).

### Statistics

Welch two-sample *t*-tests were performed using the *t.test()* function from the *stats* R package (R Core Team, 2013). Quality control to test whether the number of valid interactions was correlated to the genome size was performed by a Phylogenetics Generalized Least Square (PGLS) analysis, using *ape* (*v5.8-1I*; Paradis and Schliep, 2019) and *nlme* (*v3.1-168*; Pinheiro *et al.,* 2025) R packages (Supplemental Figure 4A). Correlations between genetic features and genomic interactions were done as for the quality control, by PGLS analysis (Supplemental Figure 4). Correlations of 3D model categories and genetic features were tested by achieving phylogenetic ANOVAs using the *phylANOVA* function with 10000 simulations from the *phytools* R package (Supplemental Figure 4; Revell, 2011).

### Visualization

The tree was converted to simmap and visualized using the *plotSimmap* function of *phytools* (*v2.5-2*; Revell, 2011) in R (Figure 1A). 3D models were visualized by loading the PDB file format from the 3DGB pipeline into Mol* Viewer (Figure 1B; Supplemental Figure 2; Sehnal *et al.,* 2021). Diagnostic plots were plotted using *hicCorrectMatrix* with the *diagnostic_plot* option from HiCExplorer, with modifications in the script for better visualization (Supplemental Figure 1). Corrected iced contact maps were plotted using *hicPlotMatrix* at 50-kb resolution with the *--vMax* option set at 2500 and the *--log1p* option from HiCExplorer (Supplemental Figure 2). All other figures were produced using the *ggplot2* package (Wickham, 2016) in R.

## Declarations

### Author contributions

AL, CC, AVM and DC conceptualized the study. AL collected data and led the methodology and software parts with contributions from all co-authors. AL and TB curated data. Formal analyses were performed by AL, CC and AVM. Data interpretation was made by AL, CC and DC. AL made data visualizations with contributions from DC. DC supervised the project. AL wrote the manuscript draft. All authors reviewed and approved the manuscript.

### Data availability

All public sequencing data used in this study are available from the NCBI Sequence Read Archive or from the Darwin Tree of Life Data Portal. Individual accession numbers and assemblies can be retrieved from the Supplemental Table 1. The phylogenetic trees, videos of rotating 3D models, genome annotations and corrected iced 50 kb contact maps generated for this manuscript are available on Zenodo, DOI: 10.5281/zenodo.17672270, 10.5281/zenodo.19015207 and 10.5281/zenodo.18982314.

### Code availability

Analysis scripts generated for this manuscript are shared on the GitHub repository https://github.com/allaigle/comparative3D and will be linked to Zenodo after acceptance of the manuscript.

## Acknowledgments

We are grateful to all the researchers from the Darwin Tree of Life project who collected samples, generated the Hi-C data and assembled the genomes used in this study. We thank Ursula Oggenfuss, Pilar Junier, Sarah Semeraro-Miéville and Alice Feurtey for advice on the manuscript.

## References

Baril T, Galbraith J, Hayward A. 2024. Earl Grey: A Fully Automated User-Friendly Transposable Element Annotation and Analysis Pipeline. Mol Biol Evol. 41(4):msae068. 10.1093/molbev/msae068

Belton J-M et al. 2015. The Conformation of Yeast Chromosome III Is Mating Type Dependent and Controlled by the Recombination Enhancer. Cell Reports. 13(9):1855–1867. 10.1016/j.celrep.2015.10.063

Bonev B, Cavalli G. 2016. Organization and function of the 3D genome. Nat Rev Genet. 17(11):661–678. 10.1038/nrg.2016.112

Boveri T. 1888. Zellenstudien II. Die Befruchtung und Teilung des Eies von Ascaris megalocephala. Jena Z. Naturwiss.

Brůna T et al. 2021. BRAKER2: automatic eukaryotic genome annotation with GeneMark-EP+ and AUGUSTUS supported by a protein database. NAR Genom Bioinform. 3(1):lqaa108. 10.1093/nargab/lqaa108

Brůna T, Lomsadze A, Borodovsky M. 2020. GeneMark-EP+: eukaryotic gene prediction with self-training in the space of genes and proteins. NAR Genom Bioinform. 2(2):lqaa026. 10.1093/nargab/lqaa026

Buchfink B, Xie C, Huson DH. 2015. Fast and sensitive protein alignment using DIAMOND. Nat Methods. 12(1):59–60. 10.1038/nmeth.3176

Burrack LS et al. 2016. Neocentromeres Provide Chromosome Segregation Accuracy and Centromere Clustering to Multiple Loci along a Candida albicans Chromosome. PLOS Genetics. 12(9):e1006317. 10.1371/journal.pgen.1006317

Camerino M, Chang W, Cvekl A. 2024. Analysis of long-range chromatin contacts, compartments and looping between mouse embryonic stem cells, lens epithelium and lens fibers. Epigenetics & Chromatin. 17(1):10. 10.1186/s13072-024-00533-x

Concia L et al. 2020. Wheat chromatin architecture is organized in genome territories and transcription factories. Genome Biol. 21(1):104. 10.1186/s13059-020-01998-1

Cremer T, Cremer C. 2001. Chromosome territories, nuclear architecture and gene regulation in mammalian cells. Nat Rev Genet. 2(4):292–301. 10.1038/35066075

Danecek P et al. 2011. The variant call format and VCFtools. Bioinformatics. 27(15):2156– 2158. 10.1093/bioinformatics/btr330

Dixon JR et al. 2015. Chromatin architecture reorganization during stem cell differentiation. Nature. 518(7539):331–336. 10.1038/nature14222

Duan Z et al. 2010. A three-dimensional model of the yeast genome. Nature. 465(7296):363–367. 10.1038/nature08973

Elliott TA, Gregory TR. 2015. Do larger genomes contain more diverse transposable elements? BMC Evol Biol. 15(1):69. 10.1186/s12862-015-0339-8

Fransz P et al. 2002. Interphase chromosomes in Arabidopsis are organized as well defined chromocenters from which euchromatin loops emanate. Proceedings of the National Academy of Sciences. 99(22):14584–14589. 10.1073/pnas.212325299

Galazka JM et al. 2016. Neurospora chromosomes are organized by blocks of importin alpha-dependent heterochromatin that are largely independent of H3K9me3. Genome Res. 26(8):1069–1080. 10.1101/gr.203182.115

Gao Q et al. 2019. Variations in Nuclear Number and Size in Vegetative Hyphae of the Edible Mushroom Lentinula edodes. Front Microbiol. 10 [accessed 2025 Oct 24]. https://www.frontiersin.org/journals/microbiology/articles/10.3389/fmicb.2019.01987/full. 10.3389/fmicb.2019.01987

Glavincheska I, Lorrain C. 2025. Three-dimensional genome architecture connects chromatin structure and function in a major wheat pathogen. BMC Biol. 23(1):353. 10.1186/s12915-025-02461-y

Gotoh O. 2008. A space-efficient and accurate method for mapping and aligning cDNA sequences onto genomic sequence. Nucleic Acids Res. 36(8):2630–2638. 10.1093/nar/gkn105

Gurevich A, Saveliev V, Vyahhi N, Tesler G. 2013. QUAST: quality assessment tool for genome assemblies. Bioinformatics. 29(8):1072–1075. 10.1093/bioinformatics/btt086

Hibbett D, Nagy LG, Nilsson RH. 2025. Fungal diversity, evolution, and classification. Current Biology. 35(11):R463–R469. 10.1016/j.cub.2025.01.053

Hoang DT et al. 2018. UFBoot2: Improving the Ultrafast Bootstrap Approximation. Molecular Biology and Evolution. 35(2):518–522. 10.1093/molbev/msx281

Hoencamp C et al. 2021. 3D genomics across the tree of life reveals condensin II as a determinant of architecture type. Science. 372(6545):984–989. 10.1126/science.abe2218

Hoff KJ et al. 2016. BRAKER1: Unsupervised RNA-Seq-Based Genome Annotation with GeneMark-ET and AUGUSTUS. Bioinformatics. 32(5):767–769. 10.1093/bioinformatics/btv661

Hoff KJ, Lomsadze A, Borodovsky M, Stanke M. 2019. Whole-Genome Annotation with BRAKER. In: Kollmar M, editor. Gene Prediction: Methods and Protocols. Springer; p 65–95 [accessed 2025 Sept 19]. https://doi.org/10.1007/978-1-4939-9173-0_5. 10.1007/978-1-4939-9173-0_5

Hsieh T-HS et al. 2020. Resolving the 3D Landscape of Transcription-Linked Mammalian Chromatin Folding. Molecular Cell. 78(3):539–553.e8. 10.1016/j.molcel.2020.03.002

Huang Y et al. 2023. HSFA1a modulates plant heat stress responses and alters the 3D chromatin organization of enhancer-promoter interactions. Nat Commun. 14(1):469. 10.1038/s41467-023-36227-3

Iwata H, Gotoh O. 2012. Benchmarking spliced alignment programs including Spaln2, an extended version of Spaln that incorporates additional species-specific features. Nucleic Acids Res. 40(20):e161. 10.1093/nar/gks708

Kalyaanamoorthy S et al. 2017. ModelFinder: fast model selection for accurate phylogenetic estimates. Nature Methods. (14):587–589. 10.1038/nmeth.4285

Katoh K, Standley DM. 2013. MAFFT Multiple Sequence Alignment Software Version 7: Improvements in Performance and Usability. Mol Biol Evol. 30(4):772–780. 10.1093/molbev/mst010

Kumar Suresh et al. 2021. Understanding 3D Genome Organization and Its Effect on Transcriptional Gene Regulation Under Environmental Stress in Plant: A Chromatin Perspective. Front Cell Dev Biol. 9 [accessed 2024 June 3]. https://www.frontiersin.org/articles/10.3389/fcell.2021.774719. 10.3389/fcell.2021.774719

Kuznetsov D et al. 2023. OrthoDB v11: annotation of orthologs in the widest sampling of organismal diversity. Nucleic Acids Res. 51(D1):D445–D451. 10.1093/nar/gkac998

Langmead B, Salzberg SL. 2012. Fast gapped-read alignment with Bowtie 2. Nat Methods. 9(4):357–359. 10.1038/nmeth.1923

Li D et al. 2022. Comparative 3D genome architecture in vertebrates. BMC Biol. 20(1):99. 10.1186/s12915-022-01301-7

Li H. 2012. seqtk Toolkit for processing sequences in FASTA/Q formats. https://github.com/lh3/seqtk

Li R et al. 2018. 3D genome and its disorganization in diseases. Cell Biol Toxicol. 34(5):351–365. 10.1007/s10565-018-9430-4

Li W, Godzik A. 2006. Cd-hit: a fast program for clustering and comparing large sets of protein or nucleotide sequences. Bioinformatics. 22(13):1658–1659. 10.1093/bioinformatics/btl158

Lieberman-Aiden E et al. 2009. Comprehensive Mapping of Long-Range Interactions Reveals Folding Principles of the Human Genome. Science. 326(5950):289–293. 10.1126/science.1181369

Lomsadze A, Ter-Hovhannisyan V, Chernoff YO, Borodovsky M. 2005. Gene identification in novel eukaryotic genomes by self-training algorithm. Nucleic Acids Res. 33(20):6494–6506. 10.1093/nar/gki937

Manni M et al. 2021. BUSCO Update: Novel and Streamlined Workflows along with Broader and Deeper Phylogenetic Coverage for Scoring of Eukaryotic, Prokaryotic, and Viral Genomes. Mol Biol Evol. 38(10):4647–4654. 10.1093/molbev/msab199

Marie P et al. 2024. Gene-to-gene coordinated regulation of transcription and alternative splicing by 3D chromatin remodeling upon NF-κB activation. Nucleic Acids Research. 52(4):1527–1543. 10.1093/nar/gkae015

Minh BQ et al. 2020. IQ-TREE 2: New Models and Efficient Methods for Phylogenetic Inference in the Genomic Era. Mol Biol Evol. 37(5):1530–1534. 10.1093/molbev/msaa015

Muszewska A, Steczkiewicz K, Stepniewska-Dziubinska M, Ginalski K. 2017. Cut-and-Paste Transposons in Fungi with Diverse Lifestyles. Genome Biol Evol. 9(12):3463– 3477. 10.1093/gbe/evx261

Neph S et al. 2012. BEDOPS: high-performance genomic feature operations. Bioinformatics. 28(14):1919–1920. 10.1093/bioinformatics/bts277

Oggenfuss U et al. 2021. A population-level invasion by transposable elements triggers genome expansion in a fungal pathogen Weigel D, Mirouze M, Joly-Lopez Z, Quadrana L, editors. eLife. 10:e69249. 10.7554/eLife.69249

O’Leary NA et al. 2024. Exploring and retrieving sequence and metadata for species across the tree of life with NCBI Datasets. Sci Data. 11(1):732. 10.1038/s41597-024-03571-y

Papantonis A, Oudelaar AM. 2025. Mechanisms of Enhancer-Mediated Gene Activation in the Context of the 3D Genome. Annual Review of Genomics and Human Genetics. 26(Volume 26, 2025):163–188. 10.1146/annurev-genom-120423-012301

Paradis E, Claude J, Strimmer K. 2004. APE: Analyses of Phylogenetics and Evolution in R language. Bioinformatics. 20(2):289–290. 10.1093/bioinformatics/btg412

Paulsen J et al. 2019. Long-range interactions between topologically associating domains shape the four-dimensional genome during differentiation. Nat Genet. 51(5):835–843. 10.1038/s41588-019-0392-0

Pinheiro J, Bates D, R Core Team. 2025. nlme: Linear and Nonlinear Mixed Effects Models. [accessed 2025 Oct 10]. https://cran.r-project.org/web/packages/nlme/index.html

Poinsignon T et al. 2023. 3D models of fungal chromosomes to enhance visual integration of omics data. NAR Genom Bioinform. 5(4):lqad104. 10.1093/nargab/lqad104

Quinlan AR, Hall IM. 2010. BEDTools: a flexible suite of utilities for comparing genomic features. Bioinformatics. 26(6):841–842. 10.1093/bioinformatics/btq033

Rabl C. 1885. Über Zelltheilung. Vol Gegenbaur C (ed) Morphologisches Jahrbuch.

Ramírez F et al. 2018. High-resolution TADs reveal DNA sequences underlying genome organization in flies. Nat Commun. 9(1):189. 10.1038/s41467-017-02525-w

Revell LJ. 2012. phytools: an R package for phylogenetic comparative biology (and other things). Methods in Ecology and Evolution. 3(2):217–223. 10.1111/j.2041-210X.2011.00169.x

Sehnal D et al. 2021. Mol* Viewer: modern web app for 3D visualization and analysis of large biomolecular structures. Nucleic Acids Res. 49(W1):W431–W437. 10.1093/nar/gkab314

Servant N et al. 2015. HiC-Pro: an optimized and flexible pipeline for Hi-C data processing. Genome Biol. 16(1):259. 10.1186/s13059-015-0831-x

Shen W, Sipos B, Zhao L. 2024. SeqKit2: A Swiss army knife for sequence and alignment processing. iMeta. 3(3):e191. 10.1002/imt2.191

Sherry S et al. 2008. NCBI SRA Toolkit Technology for Next Generation Sequence Data. https://github.com/ncbi/sra-tools

Stanke M, Diekhans M, Baertsch R, Haussler D. 2008. Using native and syntenically mapped cDNA alignments to improve de novo gene finding. Bioinformatics. 24(5):637–644. 10.1093/bioinformatics/btn013

Stanke M, Schöffmann O, Morgenstern B, Waack S. 2006. Gene prediction in eukaryotes with a generalized hidden Markov model that uses hints from external sources. BMC Bioinformatics. 7(1):62. 10.1186/1471-2105-7-62

Steenwyk JL et al. 2020. ClipKIT: A multiple sequence alignment trimming software for accurate phylogenomic inference. PLoS Biol 18(12): e3001007. 10.1371/journal.pbio.3001007

Steenwyk JL et al. 2021. PhyKIT: a broadly applicable UNIX shell toolkit for processing and analyzing phylogenomic data. Bioinformatics. 37(16):2325–2331. 10.1093/bioinformatics/btab096

Sun L et al. 2020. Heat stress-induced transposon activation correlates with 3D chromatin organization rearrangement in Arabidopsis. Nat Commun. 11(1):1886. 10.1038/s41467-020-15809-5

Tanizawa H et al. 2010. Mapping of long-range associations throughout the fission yeast genome reveals global genome organization linked to transcriptional regulation. Nucleic Acids Res. 38(22):8164–8177. 10.1093/nar/gkq955

Tavallaee G, Orouji E. 2025. Mapping the 3D genome architecture. Computational and Structural Biotechnology Journal. 27:89–101. 10.1016/j.csbj.2024.12.018

Tegenfeldt F et al. 2025. OrthoDB and BUSCO update: annotation of orthologs with wider sampling of genomes. Nucleic Acids Res. 53(D1):D516–D522. 10.1093/nar/gkae987

Ter-Hovhannisyan V, Lomsadze A, Chernoff YO, Borodovsky M. 2008. Gene prediction in novel fungal genomes using an ab initio algorithm with unsupervised training. Genome Res. 18(12):1979–1990. 10.1101/gr.081612.108

Tettey TT, Rinaldi L, Hager GL. 2023. Long-range gene regulation in hormone-dependent cancer. Nat Rev Cancer. 23(10):657–672. 10.1038/s41568-023-00603-4

[dataset] The Darwin Tree of Life Project Consortium. 2022. Sequence locally, think globally: The Darwin Tree of Life Project. Proceedings of the National Academy of Sciences. 119(4):e2115642118. 10.1073/pnas.2115642118

Tian H et al. 2020. Toward an understanding of the relation between gene regulation and 3D genome organization. Quantitative Biology. 8(4):295–311. 10.1007/s40484-020-0221-6

Wang H et al. 2024. A gap-free genome assembly of Fusarium oxysporum f. sp. conglutinans, a vascular wilt pathogen. Sci Data. 11(1):1–10. 10.1038/s41597-024-03763-6

Wickham H. 2016. ggplot2: Elegant Graphics for Data Analysis. https://ggplot2.tidyverse.org

Winter DJ et al. 2018. Repeat elements organise 3D genome structure and mediate transcription in the filamentous fungus Epichloë festucae. PLOS Genetics. 14(10):e1007467. 10.1371/journal.pgen.1007467

Wolff J et al. 2020. Galaxy HiCExplorer 3: a web server for reproducible Hi-C, capture Hi-C and single-cell Hi-C data analysis, quality control and visualization. Nucleic Acids Research. 48(W1):W177–W184. 10.1093/nar/gkaa220

[dataset] Wright R, Woof K. [accessed 2025a Oct 9]. https://wellcomeopenresearch.org/articles/8-277

[dataset] Wright R, Woof K. [accessed 2025b Oct 9]. https://wellcomeopenresearch.org/articles/8-273

[dataset] Wright R, Woof K, Gaya E. 2022. [accessed 2025 Oct 9]. https://wellcomeopenresearch.org/articles/7-83

Xia C et al. 2022. Folding Features and Dynamics of 3D Genome Architecture in Plant Fungal Pathogens. Microbiology Spectrum. 10(6):e02608–22. 10.1128/spectrum.02608-22

Yamasaki YY et al. 2025. 3D Genome Constrains Breakpoints of Inversions That Can Act as Barriers to Gene Flow in the Stickleback. Molecular Ecology. 34(21):e17814. 10.1111/mec.17814

[dataset] Yildirir G, et al. 2022. Long reads and Hi-C sequencing illuminate the two-compartment genome of the model arbuscular mycorrhizal symbiont Rhizophagus irregularis. New Phytologist. 233(3):1097–1107. 10.1111/nph.17842

Yu G. 2020. Using ggtree to Visualize Data on Tree-Like Structures. Current Protocols in Bioinformatics. 69(1):e96. 10.1002/cpbi.96

