## Supplemental Figures for "Comparative analyses reveal rapid turnover and emergence of transitory 3D genome architectures in the fungal kingdom"

Related to the article:

### Supplementary figures

**Supplemental Figure 1: 3D models and contact maps for the 55 species.** 3D models (left) are visualized using Mol*. Each bead of the 3D models represents 10 kb of a chromosome and is colored as a gradient to represent the chromosome it belongs to, from red for the longest chromosome to dark blue for the smallest one. Corrected iced matrices (right) are from HiCExplorer at 50 kb resolution. Species are displayed in phylogenetic order, as in Figure 1A.

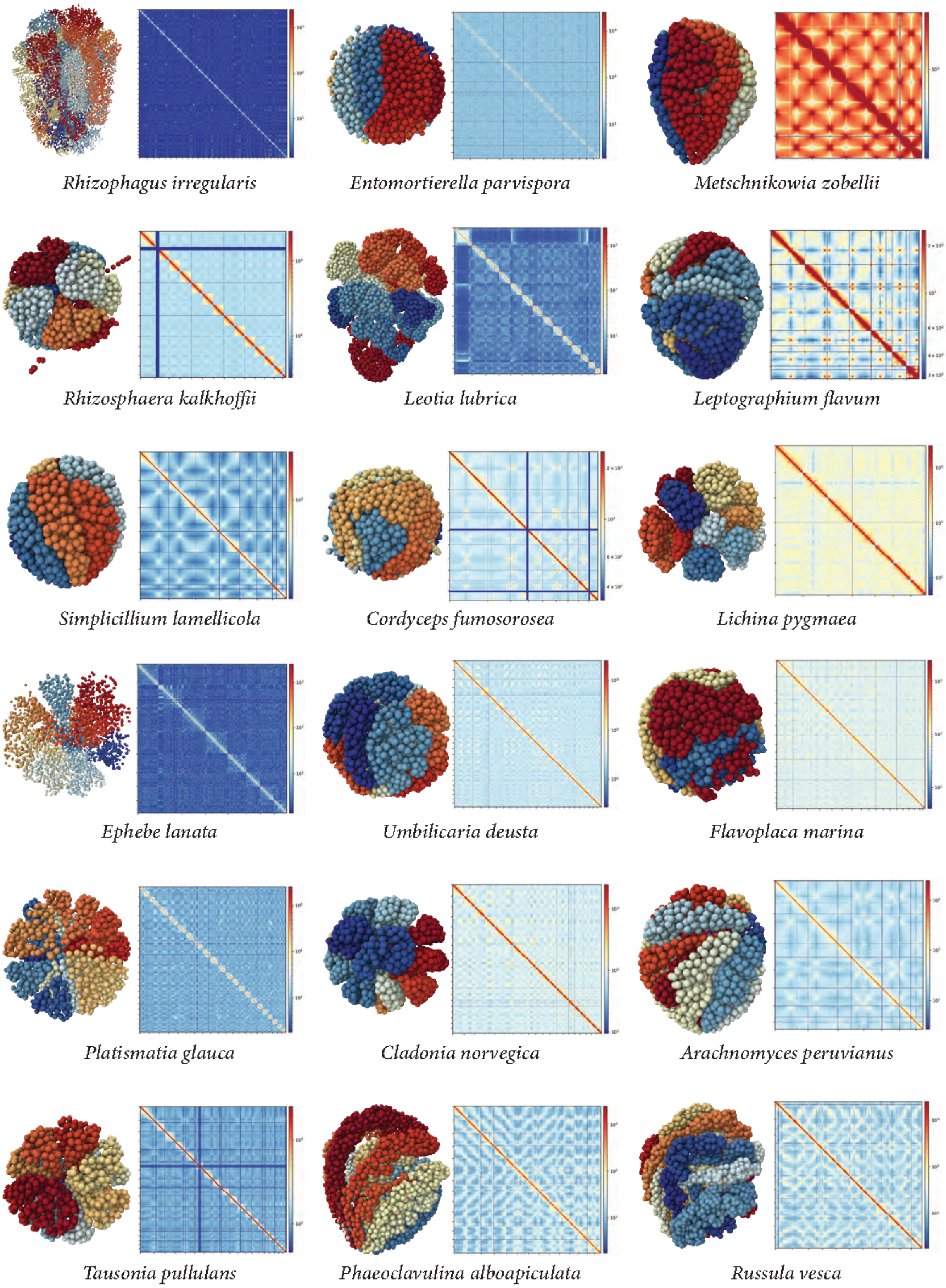
Rhizophagus irregularis
Entomortierella parvispora
Metschnikowia zobellii
Rhizosphaera kalkhoffii
Leotia lubrica
Leptographium flavum
Simplicillium lamellicola
Cordyceps fumosorosea
Lichina pygmaea
Ephebe lanata
Umbilicaria deusta
Flavoplaca marina
Platismatia glauca
Cladonia norvegica
Arachnomyces peruvianus
Tausonia pullulans
Phaeoclavulina alboapiculata
Russula vesca

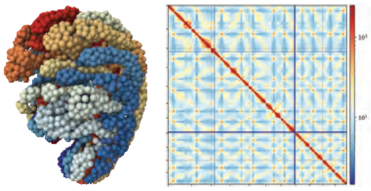
*Russula aff farinipes*

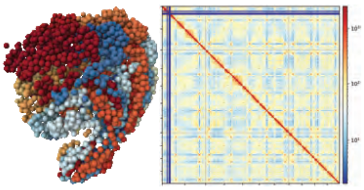
*Russula nigricans*

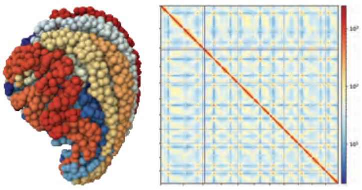
*Russula olivacea*

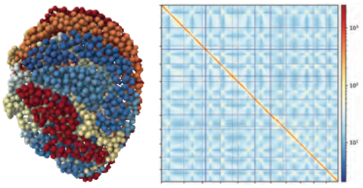
*Russula faginea*

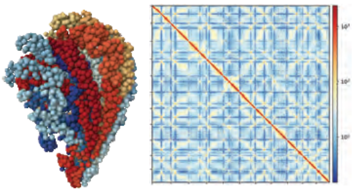
*Russula delica*

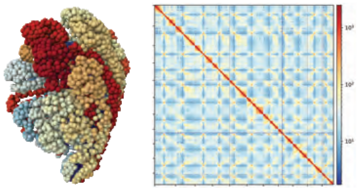
*Lactarius evosmus*

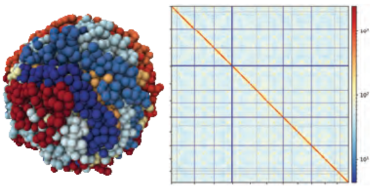
*Lactarius tabidus*

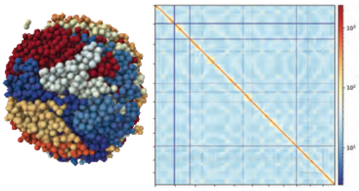
*Lactarius blennius*

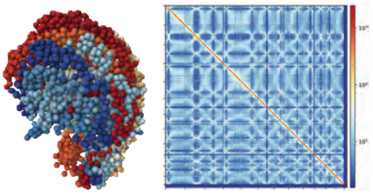
*Hericium coralloides*

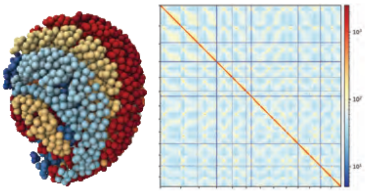
*Ganoderma resinaceum*

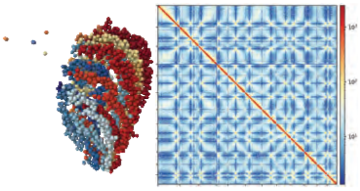
*Sparassis minoensis*

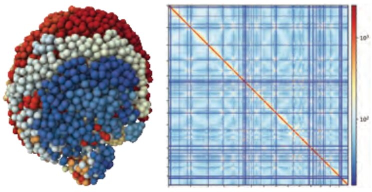
*Laetiporus sulphureus*

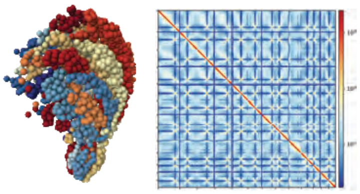
*Butyriboletus subappendiculatus*

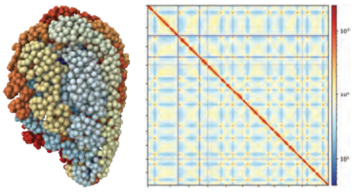
*Aureoboletus gentilis*

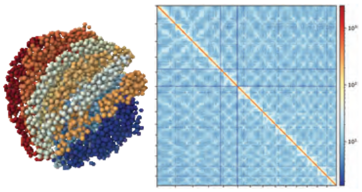
*Ramariopsis corniculata*

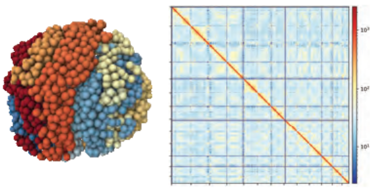
*Clavaria fumosa*

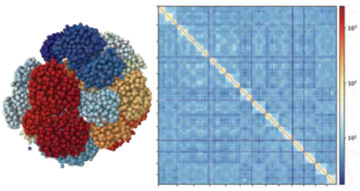
*Hygrocybe punicea*

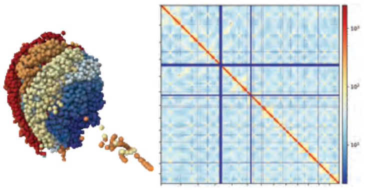
*Hygrocybe coccinea*

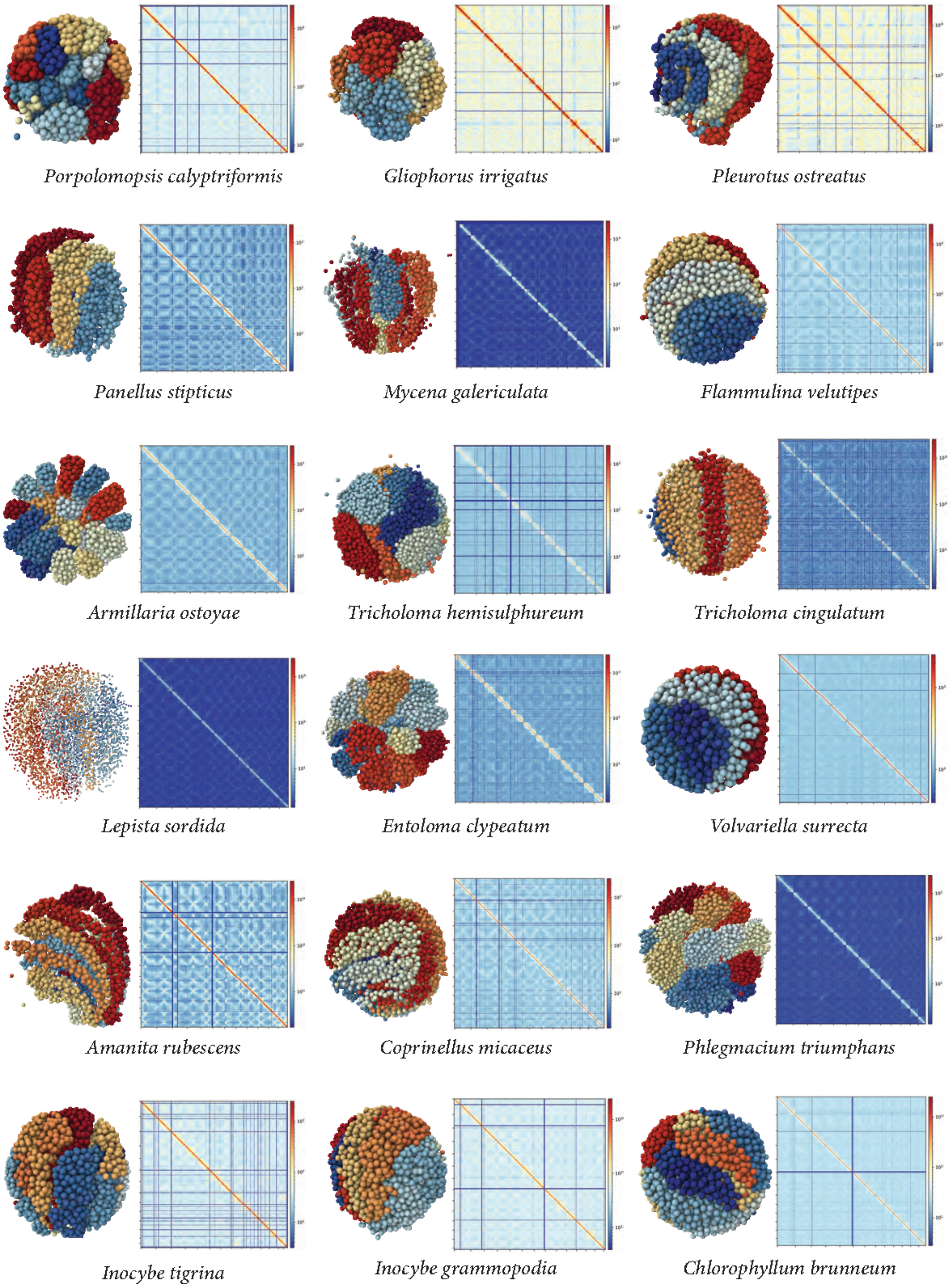
Porpolomopsis calyptriformis
Gliophorus irrigatus
Pleurotus ostreatus
Panellus stipticus
Mycena galericulata
Flammulina velutipes
Armillaria ostoyae
Tricholoma hemisulphureum
Tricholoma cingulatum
Lepista sordida
Entoloma clypeatum
Volvariella surrecta
Amanita rubescens
Coprinellus micaceus
Phlegmacium triumphans
Inocybe tigrina
Inocybe grammopodia
Chlorophyllum brunneum

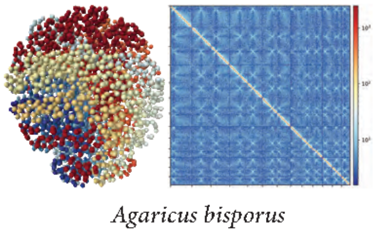


**Supplemental Figure 2: Comparison of genetic features between Ascomycota and Basidiomycota phyla. A.** Violin plot of genome size in megabase pairs by phylum (*$p$-value = 0.013; Welch Two Sample $t$-test). **B.** Violin plot of gene content in percentage by phylum (***$p$-value = 0.0021; Welch Two Sample $t$-test). **C.** Violin plot of TE content in percentage by phylum (**$p$-value = 0.0053; Welch Two Sample $t$-test).

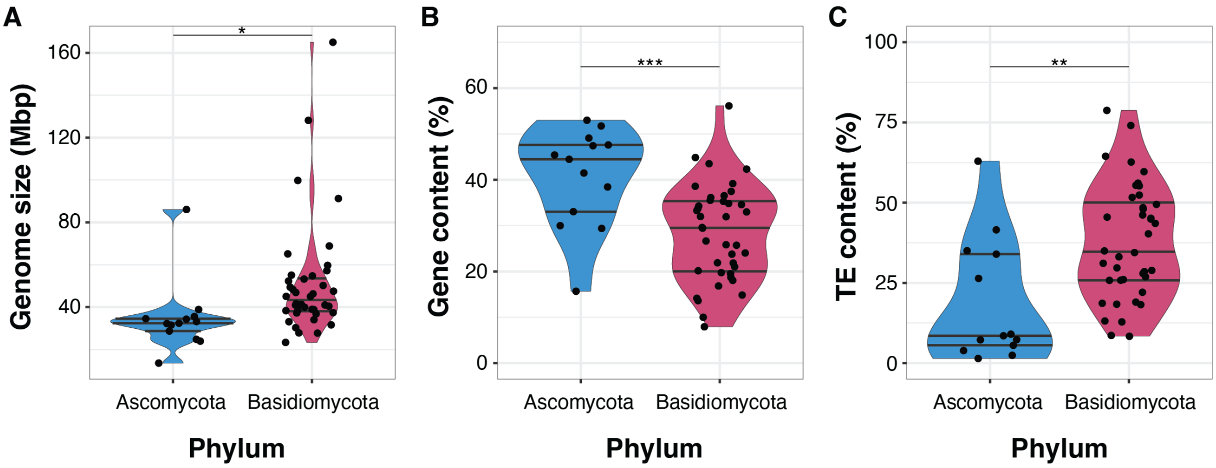


**Supplemental Figure 3: Distributions of gene and transposable elements content along the genome.** Left: schematic representation of the phylogenetic tree for the fifty-five species to distinguish their phylum. Species are phylogenetically ordered, as in Figure 2. Middle-right: Ridgeline plots of the gene and TE content in each 10 kb bin of the fifty-five genomes. The dotted black lines represent the thresholds by which we consider a region enriched in genes or in TE. For genes, over 20 % is considered gene-rich, while for TEs, under 20 % is TE-poor and over 80 % is considered TE-rich.

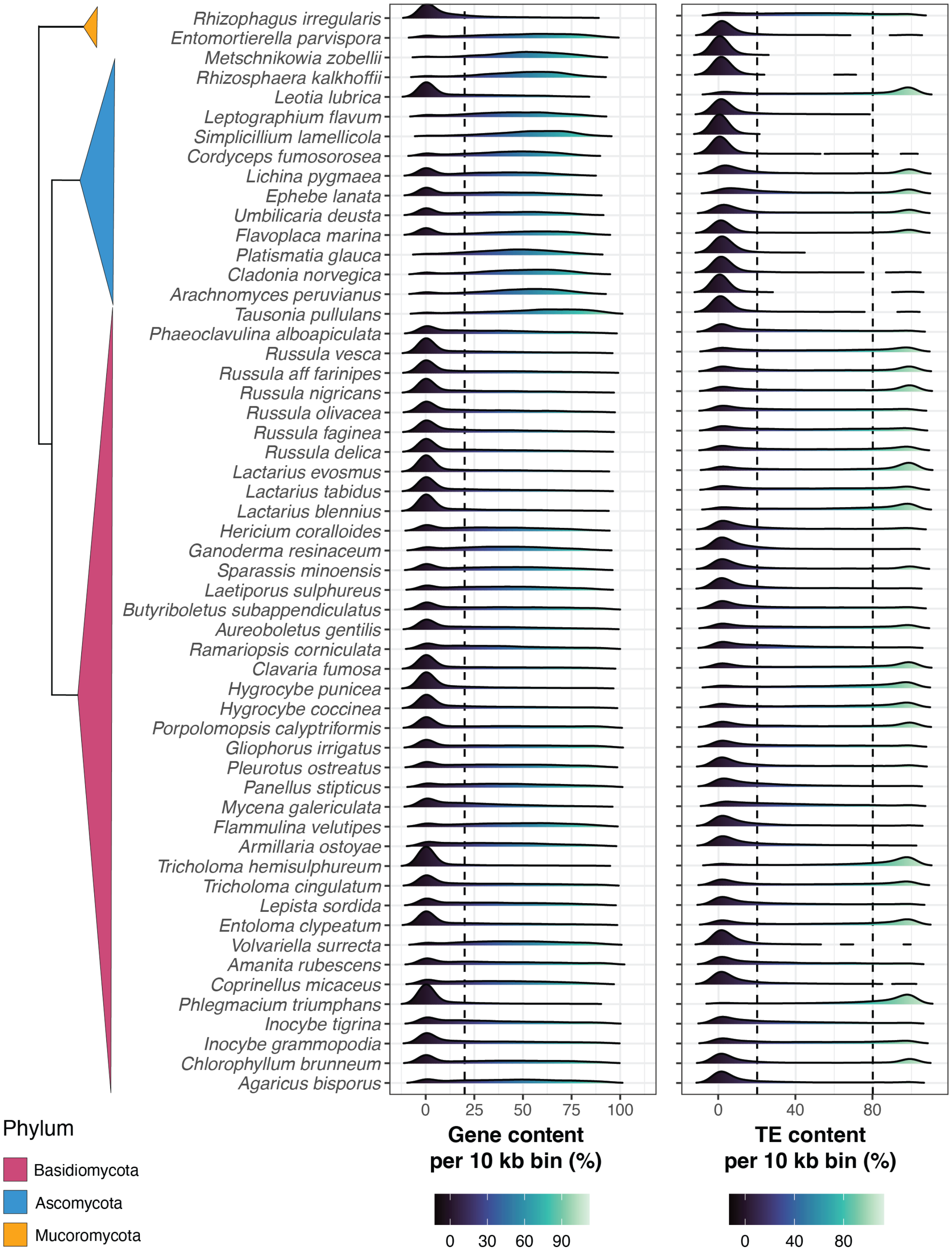


**Supplemental Figure 4: Phylogenetic regressions between genetic features across the 55 species. A-D)** Dot plots representing the presence/absence of correlation between the genome size and: **A)** total number of interactions in million (*p*-value = 0.364; AIC = 2016.869; BIC = 2024.75), B) chromosome number (**p*-value = 0.0302; AIC = 1984.436; BIC = 1992.317), **C)** gene content (%; ****p*-value = 1.9e$^{-13}$; AIC = 1940.185; BIC = 1948.066), **D)** GC content (%; ****p*-value = 3.43e$^{-05}$; AIC = 1972.277; BIC = 1980.158). **E)** Dot plot representing the presence/absence of correlation between CG content (%) and TE content (%; ****p*-value = 0.0017; AIC = 460.5116; BIC = 468.3927). **F-K)** Dot plots representing the presence/absence of correlation between the genome size and the different TE subclasses: **F)** LTR (%; *p*-value = 3.23e$^{-05}$; AIC = 1974.239; BIC = 1982.12), **G)** DNA (%; *p*-value = 8.45e$^{-09}$; AIC = 1955.44; BIC = 1963.321), **H)** LINE (%; *p*-value = 0.101; AIC = 1984.623; BIC = 1992.504), **I)** PLE (%; *p*-value = 0.756; AIC = 1981.621; BIC = 1989.502), **J)** RC (%; *p*-value = 0.101; AIC = 1985.55; BIC = 1993.431), **K)** SINE (%; *p*-value = 6.25e$^{-06}$; AIC = 1973.696; BIC = 1981.577). **L)** Dot plot representing the presence/absence of correlation between the genome size and the ratio of cis to trans interactions (%; *p*-value = 0.0377; AIC = 295.7321; BIC = 303.6132). Dot shapes and colors are as described in Figure 1.

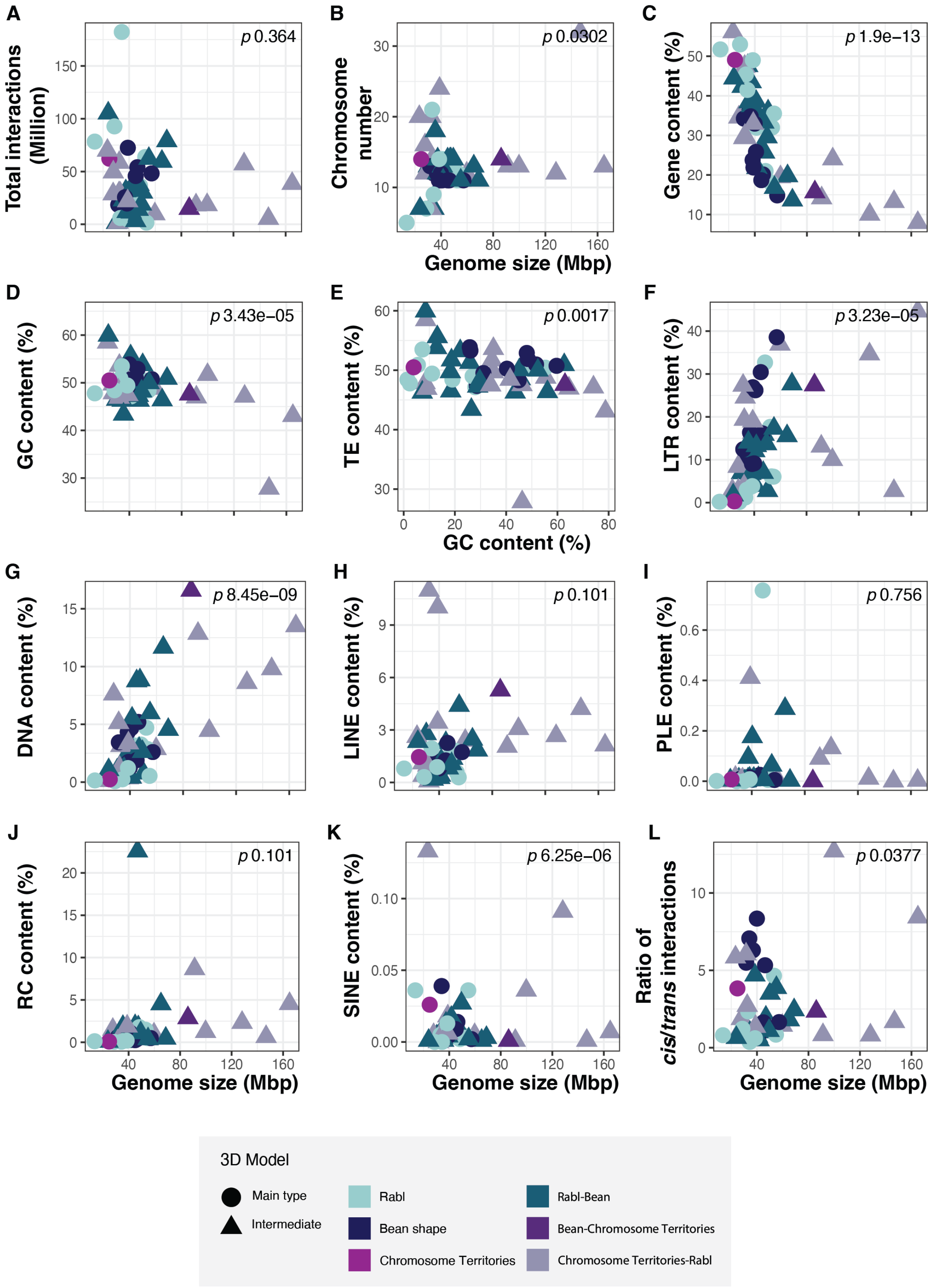
A
B
C
D
E
F
G
H
I
J
K
L
Total interactions (Million)
p 0.364
Chromosome number
p 0.0302
Gene content (%)
p 1.9e−13
Genome size (Mbp)
GC content (%)
p 3.43e−05
TE content (%)
p 0.0017
LTR content (%)
p 3.23e−05
GC content (%)
DNA content (%)
p 8.45e−09
LINE content (%)
p 0.101
PLE content (%)
p 0.756
RC content (%)
p 0.101
SINE content (%)
p 6.25e−06
Ratio of cis/trans interactions
p 0.0377
Genome size (Mbp)
Genome size (Mbp)
Genome size (Mbp)
3D Model
Main type
Intermediate
Rabl
Bean shape
Chromosome Territories
Rabl-Bean
Bean-Chromosome Territories
Chromosome Territories-Rabl

**Supplemental Figure 5: Phylogenetic ANOVAs between genetic features and 3D genome architecture across the 55 species. A-C)** Dot plots representing the presence/absence of correlation between 3D models and genetic features: **A)** the ratio of *cis* to *trans* interactions (*p*-value = 0.438), **B)** GC content (%; *p*-value = 0.7659), **C)** chromosome number (*p*-value = 0.5919) and **D)** gene content (*p*-value = 0.3062). **E-J)** Dot plots representing the presence/absence of correlation between 3D models and the different TE subclasses: **E)** LTR content (%; *p*-value = 0.1771)**, F)** DNA content (%; **p*-value = 0.0248)**, G)** LINE content (%; *p*-value = 0.1666)**, H)** PLE content (%; *p*-value = 0.9621)**, I)** RC content (%; *p*-value = 0.9166)**, J)** SINE content (%; *p*-value = 0.8003). Dot shapes and colors are as described in Figure 1.

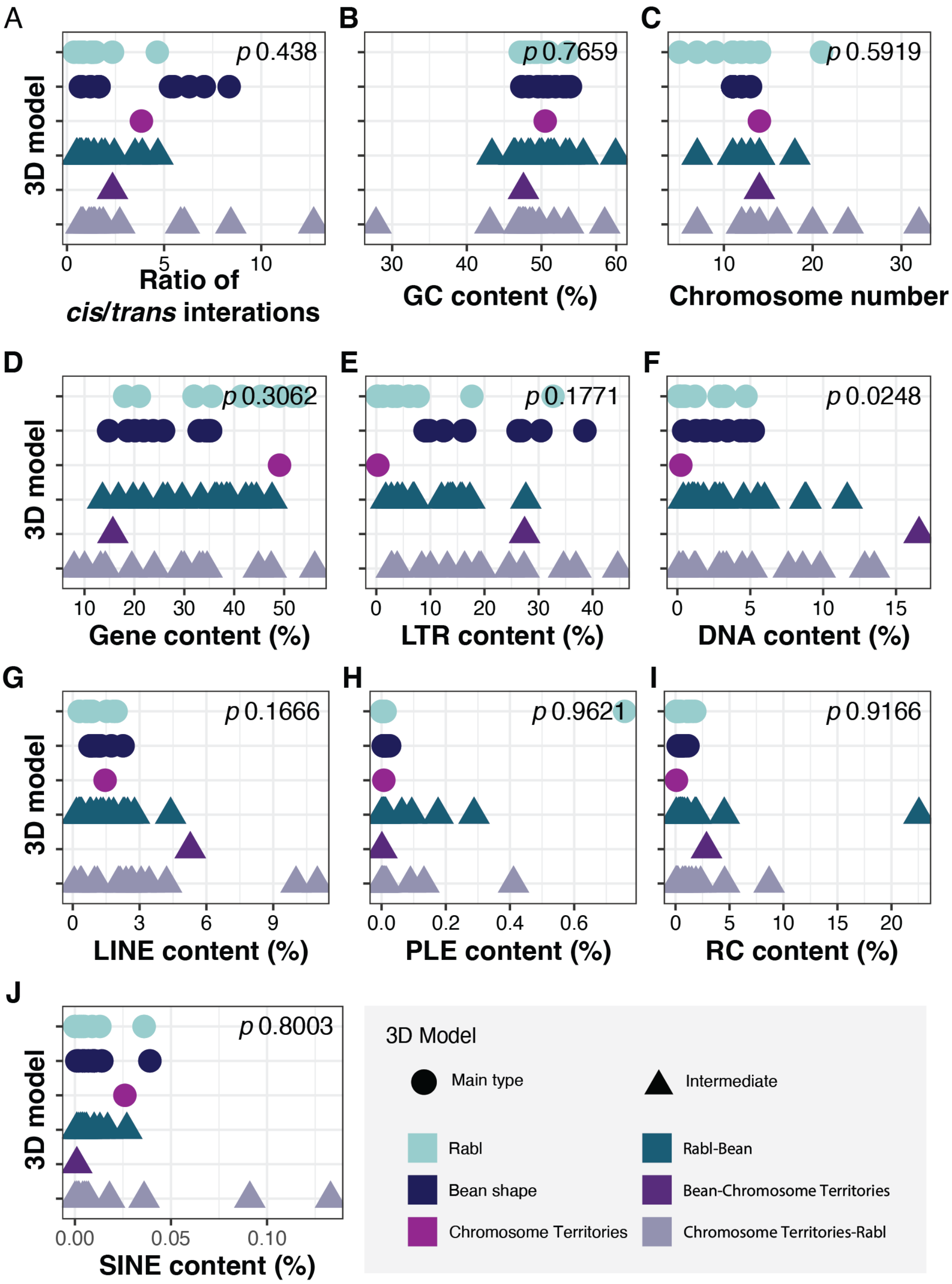


**Supplemental Figure 6: Diagnostic plots for Hi-C corrections of the 55 species.** Distribution of the frequency of interactions by the total count of bin at 50 kb resolution. The modified z-score is based on the median absolute deviation method. The blue line in each distribution plot represents the threshold chosen to correct matrices and the upper range was of 5 for all species. Species are alphabetically order.

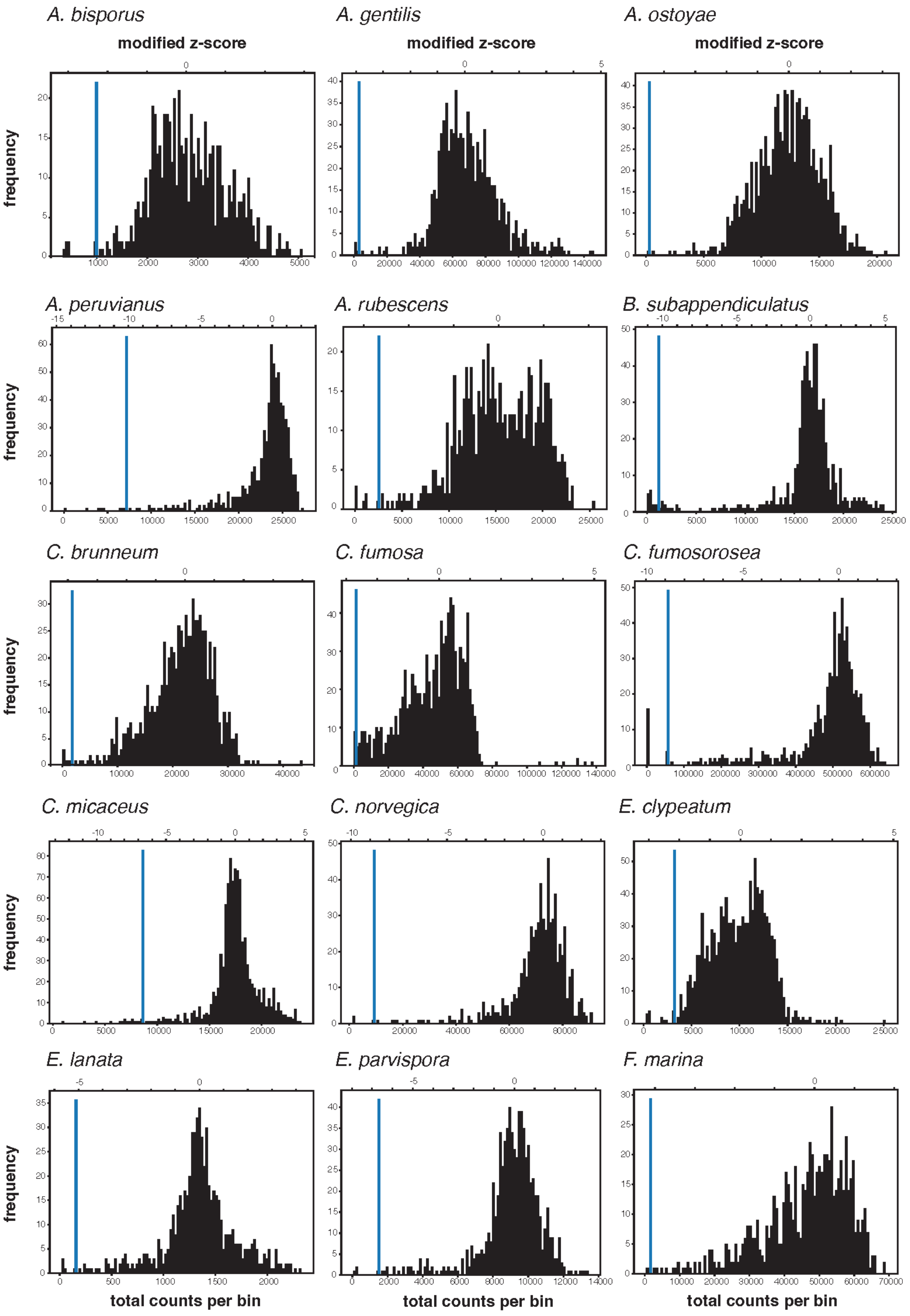
A. bisporus
A. gentilis
A. ostoyae
modified z-score
modified z-score
modified z-score
frequency
A. peruvianus
A. rubescens
B. subappendiculatus
frequency
C. brunneum
C. fumosa
C. fumosorosea
frequency
C. micaceus
C. norvegica
E. clypeatum
frequency
E. lanata
E. parvispora
F. marina
frequency
total counts per bin
total counts per bin
total counts per bin

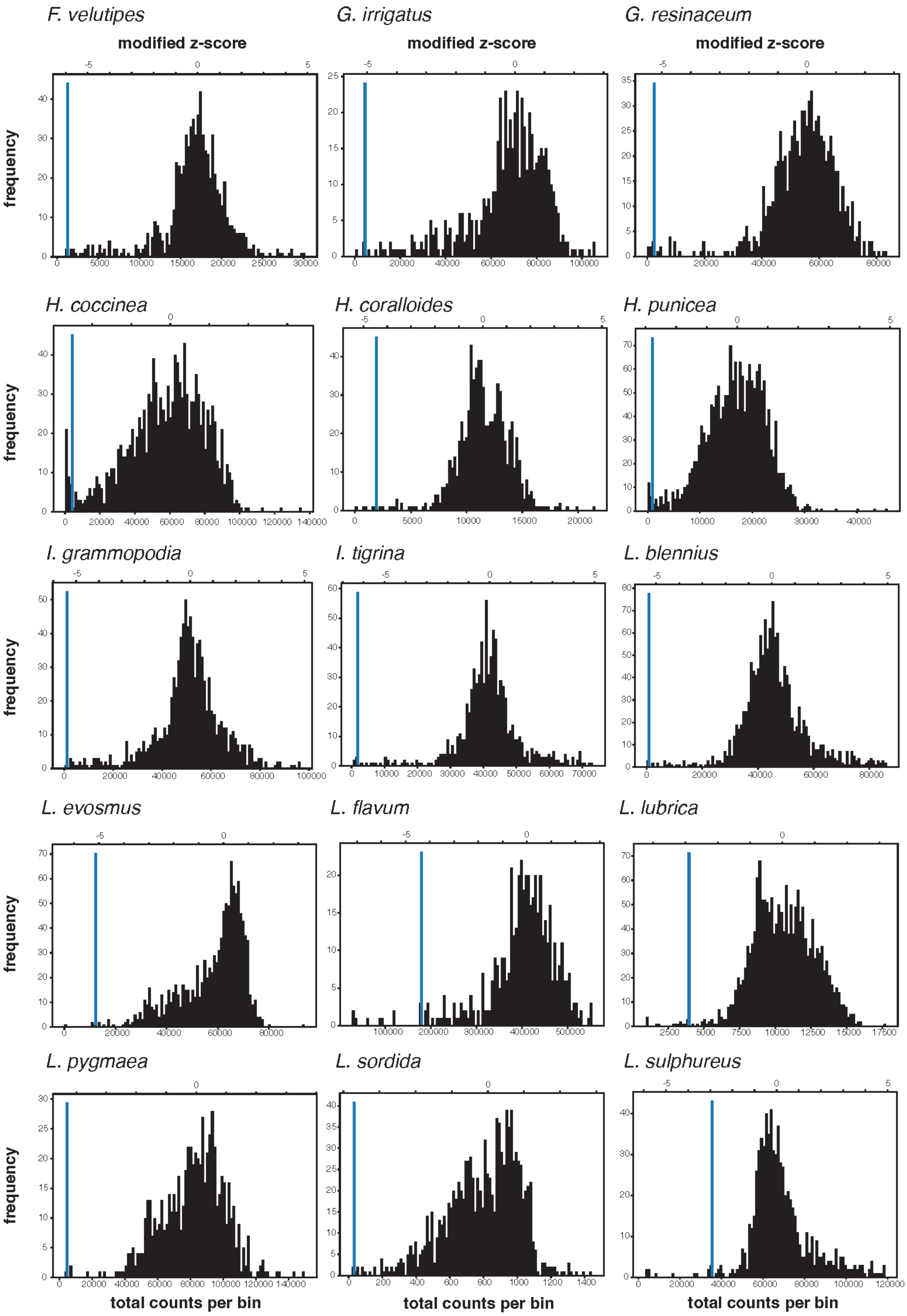
F. velutipes
G. irrigatus
G. resinaceum
modified z-score
modified z-score
modified z-score
frequency
H. coccinea
H. coralloides
H. punicea
frequency
I. grammopodia
I. tigrina
L. blennius
frequency
L. evosmus
L. flavum
L. lubrica
frequency
L. pygmaea
L. sordida
L. sulphureus
frequency
total counts per bin
total counts per bin
total counts per bin

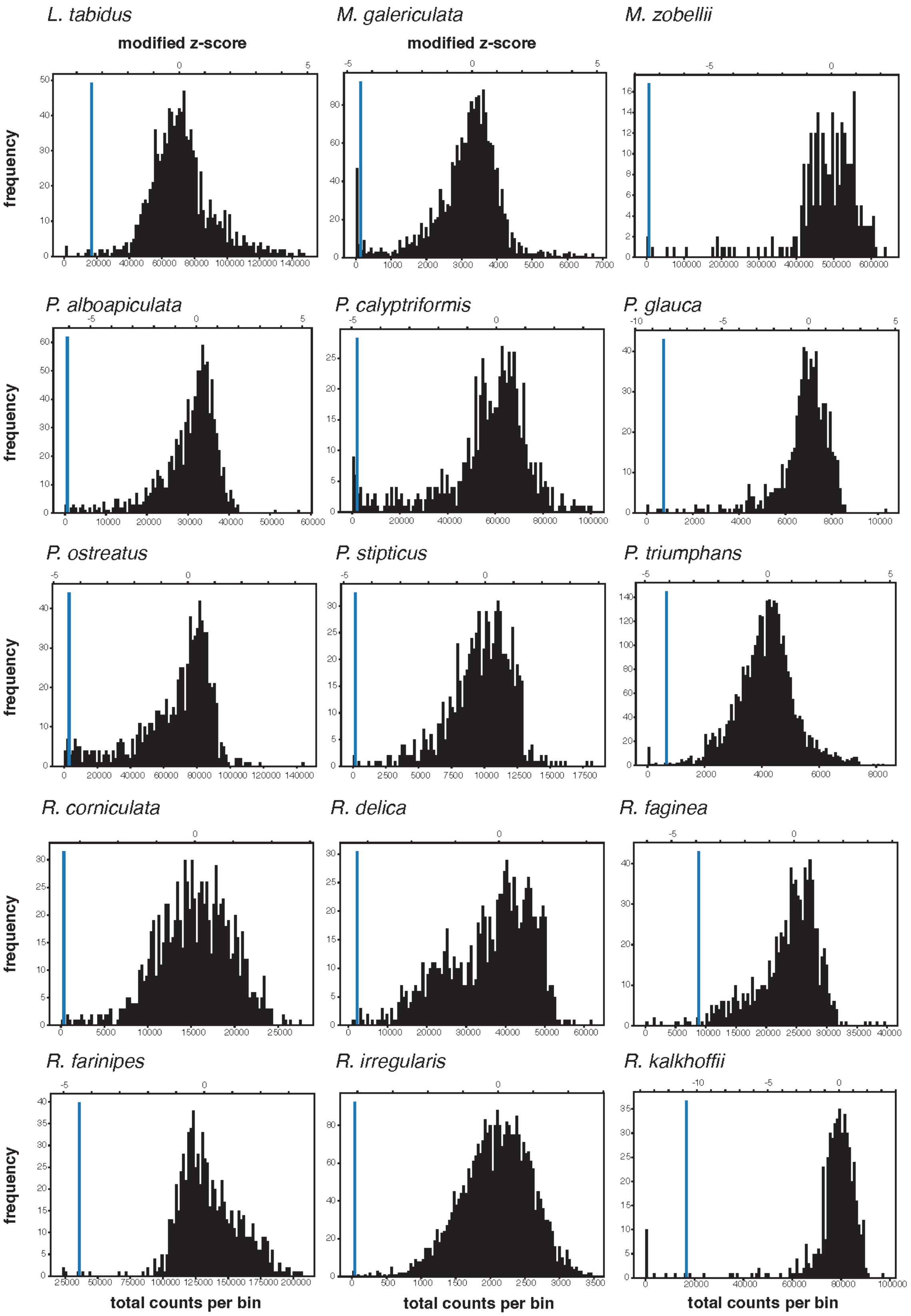
L. tabidus
M. galericulata
M. zobellii
modified z-score
modified z-score
P. alboapiculata
P. calyptriformis
P. glauca
P. ostreatus
P. stipticus
P. triumphans
R. corniculata
R. delica
R. faginea
R. farinipes
R. irregularis
R. kalkhoffii
frequency
total counts per bin
total counts per bin
total counts per bin

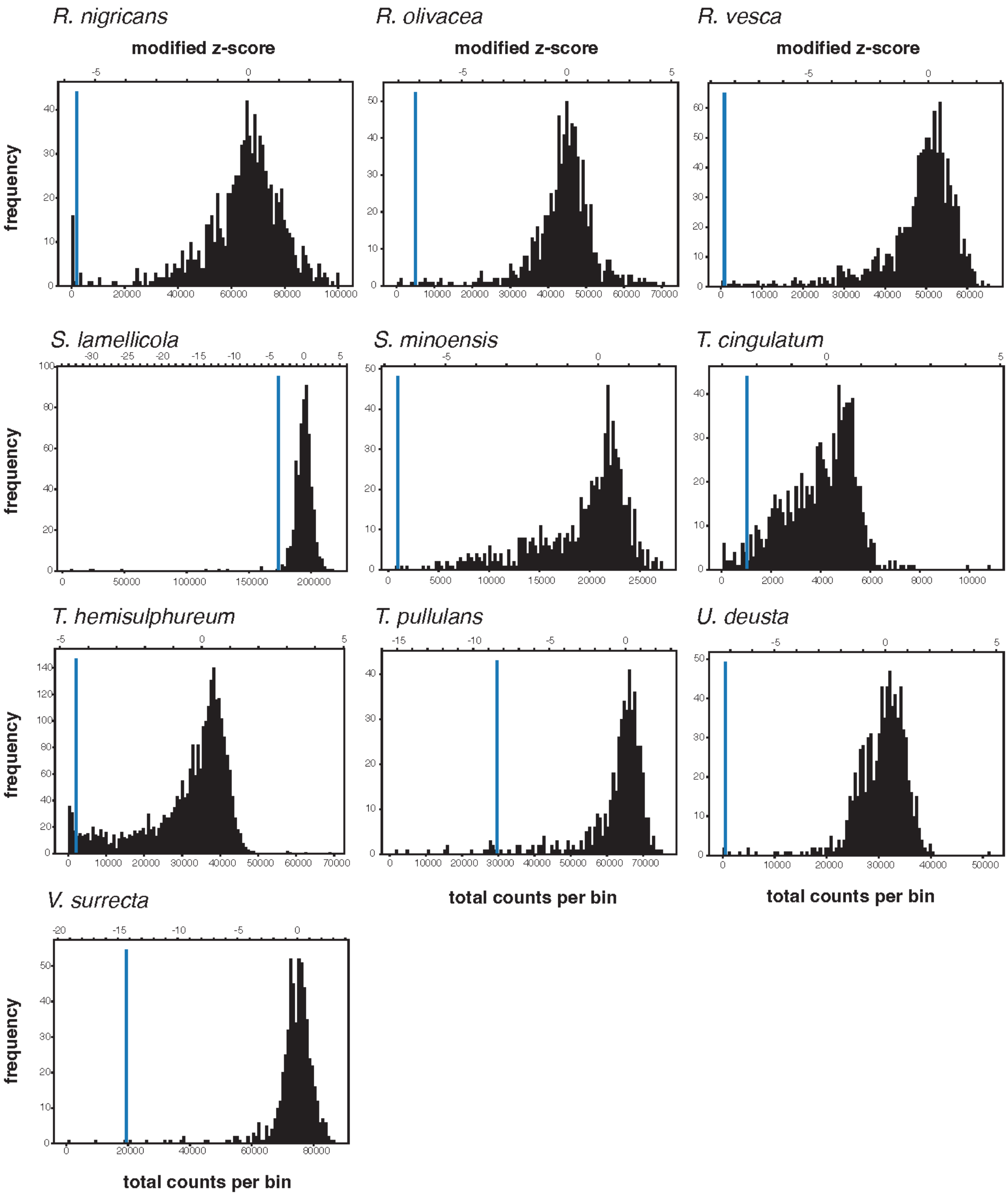
R. nigricans
R. olivacea
R. vesca
modified z-score
modified z-score
modified z-score
frequency
S. lamellicola
S. minoensis
T. cingulatum
frequency
T. hemisulphureum
T. pullulans
U. deusta
frequency
total counts per bin
total counts per bin
V. surrecta
frequency
total counts per bin
